# Multi-view graph learning for deciphering the dominant cell communication assembly of downstream functional events from single-cell RNA-seq data

**DOI:** 10.1101/2024.02.28.582416

**Authors:** Boya Ji, Xiaoqi Wang, Xiang Wang, Liwen Xu, Shaoliang Peng

**Author notes:** Corresponding authors (L. Xu); (S. Peng).

## Abstract

Cell-cell communications (CCCs) from multiple sender cells collaboratively affect downstream functional events in receiver cells, thus influencing cell phenotype and function. How to rank the importance of these CCCs and find the dominant ones in a specific downstream functional event has great significance for deciphering various physiological and pathogenic processes. To date, several computational methods have been developed to focus on the identification of cell types that communicate with enriched ligand-receptor interactions from single-cell RNA-seq (scRNA-seq) data, but to the best of our knowledge, all of them lack the ability to identify the communicating cell type pairs that play a major role in a specific downstream functional event, which we call it “dominant cell communication assembly (DCA)”. Here, we proposed scDCA, a multi-view graph learning method for deciphering DCA from scRNA-seq data. scDCA is based on a multi-view CCC network by constructing different cell type combinations at single-cell resolution. Multi-view graph convolution network was further employed to reconstruct the expression pattern of target genes or the functional states of receiver cells. The DCA was subsequently identified by interpreting the model with the attention mechanism. scDCA was verified in a real scRNA-seq cohort of advanced renal cell carcinoma, accurately deciphering the DCA that affect the expression patterns of the critical immune genes and functional states of malignant cells. Furthermore, scDCA also accurately explored the alteration in cell communication under clinical intervention by comparing the DCA for certain cytotoxic factors between patients with and without immunotherapy. scDCA is free available at: https://github.com/pengsl-lab/scDCA.git.

## 1. Introduction

Cell-cell communications (CCCs) underpin the major functions in multicellular organisms [1]. Sender cells induce cascading regulation of biological processes through CCCs, resulting in the reprogramming of the receiver cell with altered gene expression and functional pathways [2]. Depicting the landscape of CCCs not only provides insights into fundamental life processes but also offers strategies for therapeutic interventions for various complex diseases, especially cancer [3]. Numerous studies have contributed to identifying the communicating cell component pairs that act as the functional assembly inducing downstream biological processes and clinical response [4, 5]. For instance, inter-cellular communication between immune and cancer cells can modulate the expression of immune-related genes and promote tumor evasion [6]. The communication between cancer cells and cancer-associated fibroblasts (CAFs) can induce the expression of matrix metalloproteinases (MMPs), which are essential for extracellular matrix remodeling and tumor invasion [7].

Recent advances in single-cell RNA sequencing (scRNA-seq) enabled the characterization and interpretation of the complete landscape of CCC using computational methods [8, 9]. Generally, these computational methods inferred CCC through integrating scRNA-seq data and prior ligand-receptor (L-R) interaction information, and employ various models to assess the enrichment or over-presentation of L-R interactions based on the constructed background reference [10]. Two of the most widely used methods, CellPhoneDB [11] and CellChat [12], applied statistical tests to quantify the probability of each interaction over null hypothesis references. Besides, there are other methods that introduce the information of gene networks for a more accurate and complete model. For instance, NicheNet integrates the intracellular gene regulatory network [13], CytoTalk incorporated intracellular signaling networks between two cell types [14], and NATMI constructed cell-connectivity-summary networks [15]. In addition, spatial information of cells was also incorporated in some methods to refine the cell-cell interactions prediction, such as stLearn [16] and CellPhoneDB v3.0 [17].

However, to the best of our knowledge, current CCC analysis methods lack the ability to identify the communicating cell type pairs that play a major role in a specific downstream functional event. A complete characterization of CCC in microenvironment required not only the identification of communicating cell types with the enriched L-R interactions, but also their importance in affecting specific downstream functional events, e.g., the expression pattern of target genes or the functional state of recevier cells. Information exchange between different cells in microenvironment, not all cells have a direct or equal impact on downstream functional events. How to rank the importance of these CCCs and find the dominant ones has great significance for deciphering various physiological and pathogenic processes [1]. Additionaly, most of existing methods perform CCC analysis at the level of the cell type or cluster, discarding single-cell-level information. Some communication networks could be obscured with cell aggregation or downsampling [18]. In summary, it is vital to decipher CCC importance in affecting specific downstream functional events at single-cell resolution to accurately gain the full landscape of intercellular communication.

Inspired by previous researches, we noticed that the graph neural networks (GNNs) and the attention mechanism in deep learning are suitable for addressing these challenges. First, GNNs can well model the CCC networks. For example, Fischer *et al*. [19] utilized GNNs to build node-centric expression modeling (NCEM) for improving CCC inference. Yuan *et al*. [20] introduced the GCNG method, which utilized the graph convolutional neural network to infer ligand-receptor interactions. In addition, the attention mechanism in deep learning can help assess the contribution of different variables in the model and further prioritize the more important ones [21]. For example, Chen *et al*. [22] developed the GENELink method, which utilized the graph attention network to infer the high-confidence interactions between transcription factors and target genes in gene regulatory networks. A recent model called HoloNet, innovatively applied the attention mechanism to decode the functional CCC events that have a critical effect on gene expression [23].

On this basis, we defined the concept of “dominant cell communication assembly (DCA)” for the first time to denote cellular communication assemblies that dominate a specific downstream functional event. Then, we proposed scDCA, a multi-view graph learning method to decipher the DCA from single-cell RNA sequencing (scRNA-seq) data. (i) Take advantage of four state-of-the-art cell-cell communication analysis methods, scDCA systematically and reliably infers ligand–receptor (L-R) interactions from single-cell RNA sequencing data. (ii) According to prior knowledge of L-R interactions, gene expression profiles and cell type information, scDCA constructs the multi-view cell-cell communication network between different cell types at single-cell resolution by using an edge weighting strategy and filtering out edges with low specificity. (iii) scDCA develops a multi-view graph convolution network to reconstruct the expression pattern of target genes or the functional status of receiver cells and then deciphers the dominant cell communication assembly by interpreting the trained model. In a scRNA-seq cohort of advanced renal cell carcinoma, scDCA was applied to decipher the DCA that affect the expression patterns of the critical marker genes of CD8+ T cell and tumor-associated macrophages, which accurately reflected the dominant cell regulation process on these cell types. scDCA also accurately deciphered the DCA that affect different functional states of malignant cells. Additionally, scDCA was applied to explore the alteration in cell communication under clinical intervention by comparing the DCA for certain cytotoxic factors between patients with and without immunotherapy. In summary, scDCA provided a valuable and practical tool for deciphering CCCs from scRNA-seq data, which helped to reveal biologically meaningful cell communication events.

## 2. Materials and Methods

### 2.1. Data preprocessing

The scRNA-seq data of patients with advanced renal cell carcinoma were derived from the recently published study [24] and accessed via the Single Cell Portal. The entire dataset contained 34326 cells from tumor tissues of 8 patients sequenced by 10X genomics’ Visium platform. Cell samples from two patients (P76 and P915) were specifically selected. Compared to the untreated patient P76, patient P915 received immune checkpoint blockade (ICB, aPD-1 + aCTLA-4) and exhibited a clinical response. Both of them were not treated with TKI and had the same biopsy Site, histology, and tumor stage for the subsequent comparison analysis (**Supplementary Table 2**). The data preprocessing steps and cell type annotation analysis were referred to the original study. Specifically, a standard normalized unique molecular identifier (UMI) count analysis workflow, including ambient RNA-decontamination with SoupX, library-size normalization, and log-transformation with Seurat, was applied. UMAP of malignant and non-malignant cells captured across all lesions, colored by broad cell type. Granular cell types and states were discerned through iterative reprojection and unsupervised clustering of lymphoid, myeloid, and tumor compartments, and merged into broader cell type categories for visualization.

### 2.2. Inference of ligand-receptor interactions

Previously, systematic assessment and comparison analyses showed that there is a dynamics of results among different L-R interactions inference methods [10, 25]. In addition, four of the statistics-based methods showed relatively higher performance [25], they are CellPhoneDB [11], CellChat [12], NicheNet [13], and ICELLNET [26]. To obtain the more comprehensive and reliable L-R interactions between every two different cell types for the following analysis, we intend to integrate the inferred results of the above four tools. In specific, the L-R interactions supported by at least two of the four methods were retained. Take the scRNAseq data from patient P76 as a case study, we found the difference in the number of inferred L-R interactions by 4 distinct tools (NicheNet does not consider the interactions with multi-subunits, **Supplementary Fig. 1a**). Also, the overlap among the 4 methods showed the variation of the 4 methods (**Supplementary Fig. 1b**).

### 2.3. Construct multi-view CCC graphs at the single cell resolution

#### Step 1: calculation of the communication probability of a specific L-R interaction

To calculate intercellular communication probability, we modeled ligand-receptor mediated signaling interactions using the law of mass action [27]. Based on the projected scRNA-seq profiles of ligands and receptors, the communication probability *P*_*i*,*j*_ from cell *i* to *j* for a particular ligand-receptor pair *k* was modeled by:

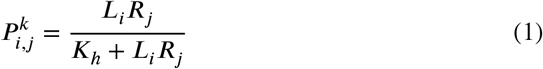

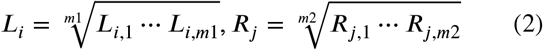

where *L*_*i*_ and *R*_*j*_ respectively represent the expression level of ligand *L* in cell i and receptor *R* in cell *j* . The exession level of ligand *L* with *m*1 subunits (i.e., *L*_*i*,1_,…, *L*_*i*,*m*_) is approximated by their geometric mean, implying that the zero expression of any subunit leads to an inactive ligand. Similarly, we compute the expression level of receptor *R* with *m*2 subunits. In addition, a Hill function is used to model the interactions between *L* and *R* with a parameter *K*_*h*_ whose default value is set to be 0.5 as the input data has a normalized range from 0 to 1.

#### Step 2: filtering out L-R interactions with low specificities

Considering the characteristic of graph learning, edges non-specifically widely expressed ligands and receptors might introduce inappropriate embeddings. To reserve the L-R interactions that actively communicating, we used a permutation test strategy to calculate the specificity of each L-R interaction. Specifically, we first randomly selected *n* background gene pairs (*n* = 200 by default) for each ligand-receptor pair. Note that the two genes of each background gene pair respectively have the most close average expression levels to the ligand and receptor. Then we calculated the communication probability of the background gene pairs to generate the null distribution of each L-R interactions. Communication probability of each background gene pair *t* between cell *i* and *j* for a particular ligand-receptor pair is:

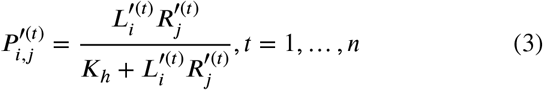

where 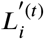 and 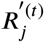 respectively represent the expression level of *t*-th background ligand and receptor in cell *i* and *j*. The communication probability of *n* background gene pairs 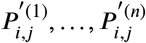 form a null distribution for the particular ligand-receptor pair. Finally, the L-R interactions whose communication probability is not over 95 percent of the null distribution will be filtered out. **Supplementary Fig. 2** showed the proportion of the altered edges in the CCC network after filtering out L-R pairs with low specificities for the P76_scRNA dataset.

#### Step 3: edge weight setting in each view of network

According to the prior knowledge, the communication probability (edge weight in the network) *w*_*i*,*j*_ between sender cell *i* and receiver cell *j* is calculated by summing up the communication probability of the filtered *n* ligand-receptor pairs between them:

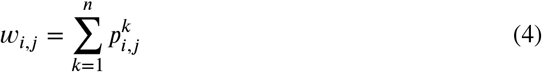

#### Step 4: construction of the entire multi-view graph

The multi-view CCC network ultimately consists of networks under different combinations of cell types, e.g. cells between cell type A and cell type B form *G*_*A*,*B*_, cells between cell type B and cell type C form *G*_*B*,*C*_, etc., where the edge weights in the network are *w*_*i*,*j*_.

### 2.4. Selection of target genes to be predicted

The selection of specific target genes was based on the following criteria: (i) they should not be genes of the mitochondrion, ligand, or receptor; (ii) they should be expressed in over 50% cells; (iii) they should be highly variable genes (detected by the *highly*_*variable*_*genes* function in Scanpy [28] package with default parameters).

### 2.5. Predicting the specific target gene expression

To distinguish the regulatory effects of CCC on gene expression and the baseline expression levels within the corresponding cell types, we hypothesized that the target gene expression *E* in all single cells could be separated into two parts:

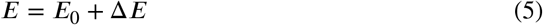

where the *E*_0_ is the baseline expression levels determined by its cell type, and Δ*E* represents the gene expression change caused by the regulatory effects of CCC. Based on this, scDCA constructed two appropriate models to reconstruct target gene expression profile.

First, a graph convolution network is constructed for predicting Δ*E*. This model can be broadly divided into 3 parts: (i) multi-view graph convolution network was applied to generate the embeddings of nodes (ii) the embeddings of nodes from each view were fused based on an attention mechanism (iii) multi-layer perceptron (MLP) model was applied to the final embeddings for predicting Δ*E*.

Specifically, the inputs to the model include the adjacency matrix *A* = {*A*^1^, *A*^2^, …, *A*^*k*^} of the multi-view communication network and the initial feature matrix *X* of the nodes (cells). The adjacency matrix *A*^*k*^ ∈ *R*^*NxN*^, where *k* represents the CCC network for the *k*-th combination of cell types, *N* represents the number of cells, and the elements in the matrix 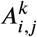 represent the edge weights (communication probability) *w*_*i*,*j*_ between cell *i* and *j*. The initial feature matrix *X* ∈ *R*^*NxM*^ is generated by the one-hot encoding method based on cell type, where *M* represents the number of cell types. To obtain the embedding of nodes in each view, the GCN model [29] used each matrix in *A* as the adjacency matrix and *X* as the initial feature matrix. Then, the final embeddings of nodes were obtained by applying the attention mechanism to integrate the embeddings from each view. Finally, the MLP was used to predict Δ*E* ∈ *R*^*N*^ :

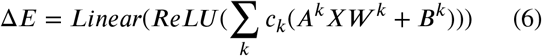

where *c*_*k*_ represents the learnable attention score, *W* ^*k*^ represents the weight matrix, *B*^*k*^ represents the bias matrix of *k*-th view, and *ReLU* is the activation function:

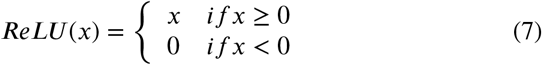

Second, the MLP model is constructed for predicting *E*_0_ based on the cell type matrix *X*:

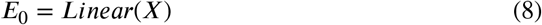

Then the above two separate models jointly predict the get gene expression *Ê* :

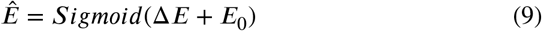

where *Sigmoid* is the activation function:

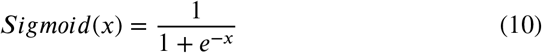

The mean squared error (MSE) between *Ê* and true expression profile *E* was used as the loss function in this work:

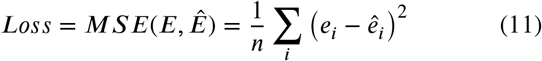

### 2.6. Calculation of the functional states activity of malignant cells

To characterize the functional states of each malignant we cell, obtained the signature gene sets of 14 crucial functional states of cancer cells, including stemness, invasion, metastasis, proliferation, epithelial-mesenchymal transition (EMT), angiogenesis, apoptosis, cell cycle, differentiation, DNA damage, DNA repair, hypoxia, inflammation and quiescence from the CancerSEA database (http://biocc.hrbmu.edu.cn/CancerSEA/) [30]. Based on these signatures, the activities of 14 functional states across malignant cells were calculated using Gene Set Variation Analysis (GSVA) with the GSVA package in R.

### 2.7. Predicting the functional state of the receiver cell

With a similar hypothesis to predicting target gene exession, the functional state activity *F* of the receiver cell was also separated into two parts:

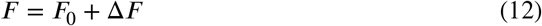

where the *F*_0_ is the baseline functional state activity for each receiver cell, and Δ*F* represents the change of functional state activity caused by the regulatory effects of CCC. Also, the scDCA integrated the graph convolution network and MLP models to predict the functional state activity of receiver cells, similar to the construction process in predicting target gene expression. The only difference is that we only consider the functional state alteration of receiver cells (malignant cells here), and then the input adjacency matrixes (views of CCC graph) were limited to the communication between every non-malignant cell type and malignant cells. Furthermore, the initial feature of GCN is randomly initialized instead of the one-hot coding.

### 2.8. Training strategy

We divide any dataset into training (85% cells) and validation sets (15% cells) of cell samples. We ran the model for 500 epochs on the training sets and chose the model with the smallest MSE on the validation sets as the final model for the subsequent interpretation analysis. The Adam optimizer (initial learning rate of 0.1 and weight decay of 5 × 10^−4^) and the StepLR learning rate decay method (step size of 10 and gamma of 0.9) was respectively used. To ensure the robustness of the model and the reliability of the interpretation, the model was repeatedly trained for 50 times. The predicted expression profile and functional state activity shown in the results are the average results obtained from each repetition.

### 2.9. Model interpretation for identifying the dominant cell communication assembly

As mentioned above, we prioritized the views based on the attention scores which represent the contribution of the corresponding cell communication assembly on downstream gene expression or cell functional state. The attention score used for prioritization was the average value of 50 repeatedly trained models. The top-ranked cell communication assembly was defined the dominant cell communication assembly. The ratios of Δ*E*/(Δ*E* + *E*_0_) and Δ*F* /(Δ*F* + *F*_0_) reflected the extent that gene expression or cell functional state is affected by cell communication. The ratio is closer to 1, the more regulatory effect from cell communication on gene expression and cell function process.

### 2.10. Collection of expression and clinical information from other independent ccRCC cohorts

Bulk RNA-seq samples of the kidney renal clear cell carcinoma (KIRC) from The Cancer Genome Atlas (TCGA) (N = 607) was obtained. The normalized FPKM data was collected through the UCSC Xena database of the GDC project (https://xena.ucsc.edu/). Another two independent ccRCC cohorts (here denoted as Motzer_NatMed_2020 and Braun_NatMed_2020_Checkmate025) with the treatment condition information were obtained from previous analysis to compare the prognostic effects of L-R pairs between immunotherapy and targeted therapy subpopulations. Normalized RNA-seq, treatment condition, and progression-free survival (PFS) data were collected from cohort NCT02684006, which incorporated with first-line avelumab + axitinib (anti-PD-L1 with tyrosine kinase inhibitor (TKI)) vs sunitinib (multitarget TKI) in advanced renal cell carcinoma [31]. Also, we obained bulk RNAseq and overall survival data of pre-treatment samples from advanced ccRCC in a randomized clinical trial (Checkmate 025) comparing the mTOR inhibitor everolimus with nivolumab (anti-PD-1) [32].

### 2.11. Statistics analysis

For comparisons of the continuous value between different groups, a two-sided Wilcoxon rank-sum test was performed. Co-expression of genes in TCGA samples was assessed based on Pearson’s correlation analysis. For the survival analysis, Kaplan-Meier survival curves were generated and the log-rank test was used to determine the significant differences among patient groups. The hazard ratio (HR) and the 95% confidence interval (CI) were calculated using univariate Cox regression analysis. All statistics analyses were performed in R version 4.2.2.

### 2.12. Functional enrichment analysis

Collectively, the g:Profiler tool (https://biit.cs.ut.ee/gprofiler/gost) was used to perform the functional enrichment analysis of Gene Ontology (GO) and the Kyoto Encyclopedia of Genes and Genomes (KEGG) pathways. The results of the enrichment analysis were visualized with the dot plot. To compare the genes more affected or less by CCC, we selected 5867 highly variable genes as targets and ranked them based on the performance improvements after considering CCC, i.e. difference of the Pearson’s correlation coefficient between the predicted expression considering CCC effects or not and the real expression level). The top-ranked and bottom-ranked 50 genes were respectively selected for GO-biological process (GO-BP) enrichment analysis.

## 3. Results

### 3.1. The overview of scDCA workflow

The goal of scDCA is to decipher the dominant cell communication assembly (DCA), i.e., CCCs with a dominant influence in a specific downstream functional event in the tumor microenvironment. scDCA requires two inputs: a gene-by-cell count matrix and a cell type label vector. The main application scenarios include the following: (1) to decipher the DCA that affect the expression pattern of target genes (2) to decipher the impact of CCC for different cell types (3) to decipher the DCA that affect the functional states of recevier cells (4) to explore the alteration of DCA in cell communication under clinical intervention. The workflow of scDCA involve four successive steps (**Fig. 1**), as briefly described below. More technical details are given in Methods.

**Figure 1:**
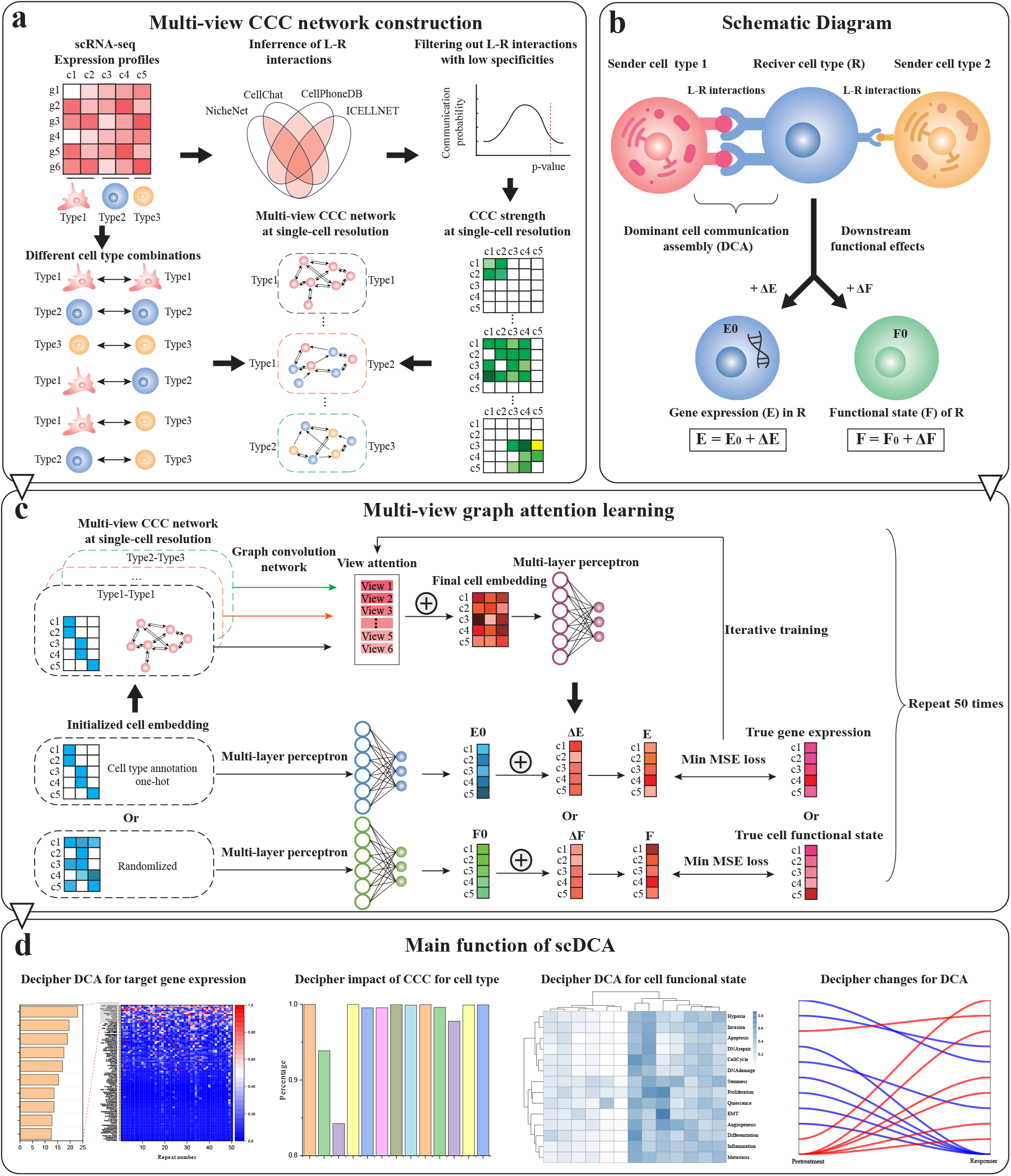
The overview of scDCA workflow. **(a)** The construction of multi-view cell-cell communication (CCC) network at single-cell resolution. See section 3.1 for details. **(b)** The schematic diagram of a full picture of CCC and its downstream functional events. LR: Ligand-Receptor; *E*: gene expression profile; *E*_0_: baseline expression determined by cell type; Δ*E*: expression change caused by CCC. *F* : functional states activity of receiver cells; *F*_0_: baseline state activity determined by the individual receiver cell; Δ*F* : change of state activity caused by CCC. **(c)** The multi-view graph convolution network for reconstructing target gene expression patterns or functional states of receiver cells. MSE: Mean Squared Error. See section 3.1 for details. **(d)** The main function of scDCA.

#### Step 1: Construction of multi-view CCC network

scDCA is based on a multi-view CCC network between different cell types at single-cell resolution (**Fig. 1.a**). First, four excellent CCC analysis tools were integrated to infer the ligand-receptor (L-R) pairs from the scRNA-seq expression profile **(see section 2.2 for details)**. Second, L-R interactions with low specificities would be filtered out and then the communication probability between a pair of single cells was calculated based on these remaining L-R interactions **(see section 2.3 for details)**. Finally, the multi-view CCC network between each pair of different cell types was constructed based on the CCC adjacency matrix, where the nodes in the graph represent individual cells and the edges represent the communication probability between them.

#### Step 2: Decomposition of gene expression and cell functional state

Briefly, cell signaling from the sender cell is transmitted to the receiver cell and elicits downstream responses, such as alteration of gene expression or even the cell states of the receiver cell. Therefore, the expression (*E*) of the specific target gene in the receiver cell can be defined as the composite of altered gene expression regulated by CCC (Δ*E*) and the baseline expression levels inherent in receiver cell type (*E*_0_). Similarly, we defined the functional state of the receiver cell (*F*) be the composite of the changed functional state caused by the regulatory effects of CCC (Δ*F*) and the baseline functional state activity for the receiver cell (*F*_0_).

#### Step 3: Reconstruction of gene expression and cell functional state

scDCA constructed two models to reconstruct the (Δ*E*/Δ*F*) and (*E*_0_/*F*_0_) (**Fig. 1.c**). For predicting the Δ*E*/Δ*F*, each view of the CCC graph was trained by a graph convolution network, node embeddings of each view were integrated through the attention mechanism and then retrained using a multi-layer perceptron (MLP). For predicting the *E*_0_/*F*_0_, another MLP was directly applied based on the cell type information or individual cell information. Finally, two estimations were summed as the final prediction results 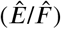 . The model was trained iteratively to minimize the mean squared error (MSE) between 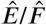 and true expression profile (*E*/*F*) **(see section 2.5 and 2.7 for details)**.

#### Step 4: Interpretation of trained model

After the model converges, these different views were prioritized based on the attention weights of scDCA which represented the contribution of the corresponding cell communication assembly on downstream functional events (**Fig. 1.c**). To ensure the reliability of the results, we repeated the training procedure and used the average attention value as results. The communicating cell type pair corresponding to the higher ranked views were regarded as the DCAs (**Fig. 1.d**).

### 3.2. The CCCs are heterogeneous at single-cell resolution

To verify the necessity of analyzing CCC at single-cell resolution, we inferred the intercellular communication status of an advanced renal cell carcinoma microenvironment dataset from a specific patient P76 **(see section 2.1)**. We inherited the cell type annotation from the data resource analysis [24], and performed the uniform manifold approximation and projection (UMAP) analysis to verify the different cell populations (**Fig. 2.a**). We found that malignant, TAM, NK, and CD8+ T cells had greater proportion (**Fig. 2.b**).

**Figure 2:**
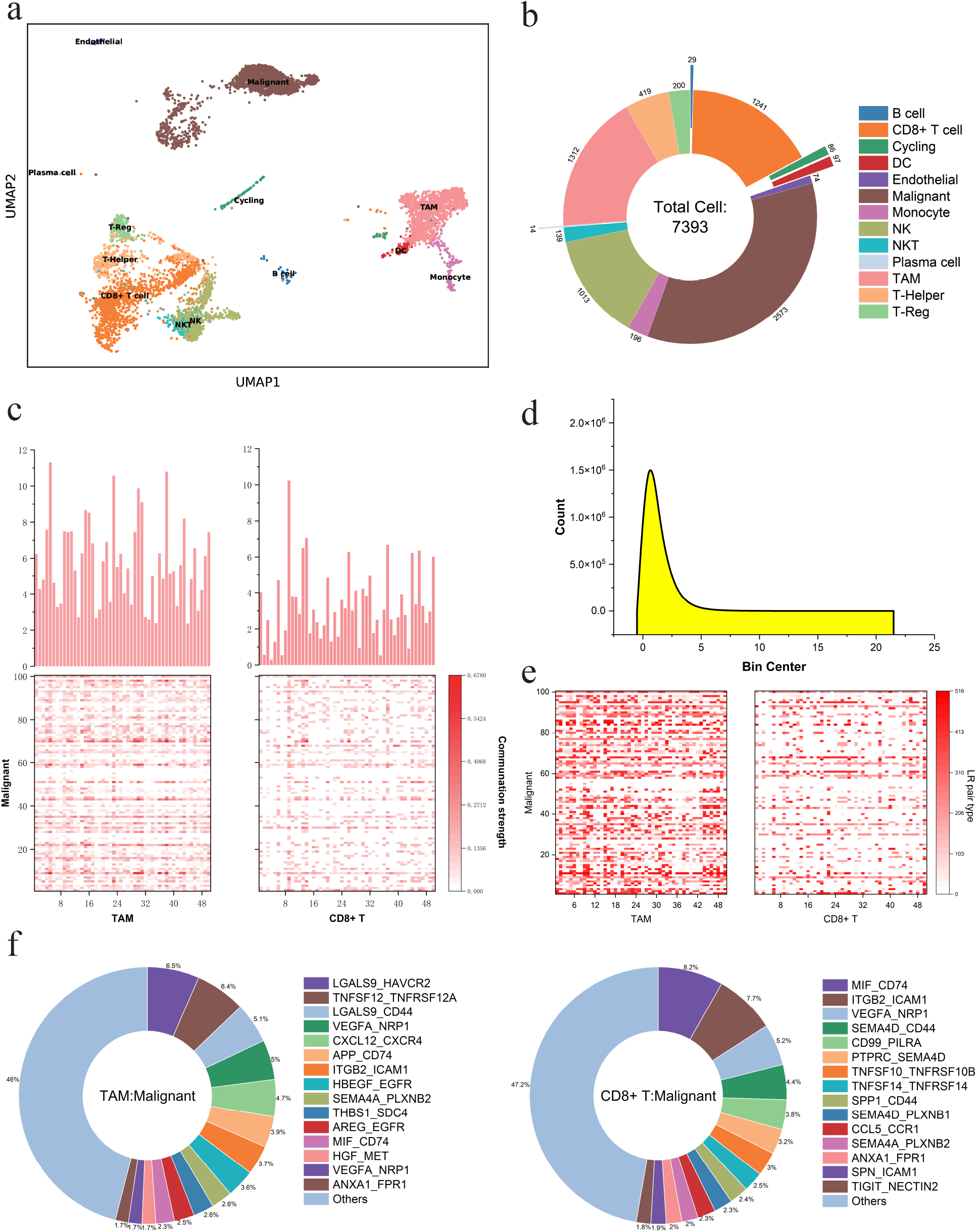
The CCCs are heterogeneous at single-cell resolution in patient P76. **(a)** UMAP of all captured cell samples from patient P76. Cell type annotation was inherited from the data resource analysis and visualized with different colors. DC, dendritic cell; NK, natural killer cell; NKT, natural killer T cell; TAM, tumor-associated macrophage; T-Reg, regulatory T cell. **(b)** The proportions of all cell types in the dataset. **(c)** Heatmap of the communication strength between every cell pair from 50 randomly selected TAMs or CD8+ T cells (column) and 100 randomly selected malignant cells (row). The histogram showed the sum of communication strength with all malignant cells for each TAM or CD8 + T cell (column), that is, the sum of each column of the heatmap. **(d)** Distribution of the number of L-R pairs in different cell pairs from **c. (e)** Heatmap showed the different types of L-R pair with maximum communication strength between every cell pair from 50 randomly selected TAMs or CD8+ T cells (column) and 100 randomly selected malignant cells (row). Color indicted different numbered L-R pairs. **(f)** Proportions of different L-R pairs mediating cellular communication of each cell pair between TAM (left) or CD8+ T cell (right) and malignant cells.

Then, we demonstrated the heterogeneity of CCC between different cells at the single-cell resolution from different perspectives. We randomly selected 100 immune cells (50 TAM cells and 50 CD8+ T cells) and 100 malignant cells, and then calculated and compared the communication strength between the two types of immune cells and malignant cells at the single-cell resolution **(see section 2.3 for details)**. It is obvious from the heatmap in **Fig. 2.c** that for the same malignant cell, different types of individual immune cells send out communication signals with different strengths (the color shades in the heatmap represents the strength of CCC between two individual cells). In addition, even for individuals of the same immune cell type, their communication strength with malignant cells varies dramatically (The histogram in **Fig. 2.c** represents the total communication strength of individual immune cells with malignant cells). **Fig. 2.d** shows the distribution of the number of communicational L-R pairs between immune and malignant cells in **Fig. 2.c**. In particular, most of the cell pairs communicated through approximately one L-R pair, very few cell pairs produce more than 4 L-R pairs-mediated CCC. Besides, we further compared the top L-R pair with the highest communication strength between immune and malignant cells in **Fig. 2.c**. The color shades in **Fig. 2.e** represent the type of the top L-R pair, again showing a high degree of heterogeneity. Finally, we counted the proportions of different L-R pairs expressed in all cell pairs between TAMs or CD8+ T cells and malignant cells. Unsurprisingly, either TAMs or CD8+ T cells exhibited diverse landscapes of L-R pairs mediating CCC with malignant cells (**Fig. 2.f**). In summary, CCCs are highly heterogeneous at the single-cell resolution. By considering CCCs at the single-cell resolution rather than at the cell type level, signals from different cellular sources can be integrated to provide a more comprehensive picture of the real CCCs and downstream functional events in the tumour microenvironment.

### 3.3. The DCA that affects the CD8A gene expression pattern is accurately deciphered

To demonstrate the ability of scDCA to accurately decipher the DCA of specific target genes, we applied scDCA to scRNA-seq data from advanced renal cell carcinoma (P76_scRNA, **see section 2.1 for details**). Specifically, we constructed a multi-view CCC network involving 13 cell types, resulting in 169 views that systematically depict the CCC in the tumor microenvironment (TME). We selected CD8A (CD8 Subunit Alpha) as a target gene example (**Fig. 3.a**). It codes for a subunit of cell surface glycoprotein CD8 commonly found on most cytotoxic T lymphocytes (CD8+ T cells), and mediates efficient CCCs involving anti-tumor immune response. As shown in **Fig. 3.b**, the reconstructed CD8A expression pattern by scDCA was highly consistent with the actual expression profile and showed a significant correlation (Pearson correlation r = 0.702, p-value *<* 0.05). After training, the ratio of Δ*E* to total expression levels (Δ*E* + *E*_0_) for each cell type was calculated, which represented the degree to which CD8A gene expression is affected by CCC in different cell types. As shown in **Fig. 3.c**, the expression of CD8A in CD8+ T and Cycling cells was less affected by CCC and more contributed by *E*_0_. This corresponds to the cell types in which CD8A is highly specifically expressed (**Fig. 3.a**). In addition, scDCA went on to decipher the expression patterns of all 5867 highly variable genes (HVGs) in the dataset, identifying genes that were either more or less affected by CCC. The difference in Pearson correlation coefficients between the gene expression profiles reconstructed by scDCA and the actual expression profiles before and after the addition of the CCC network was used as an index to rank the HVGs. Through GO enrichment analysis, the top 50 affected HVGs by CCC showed functional roles involving cell communication, cellular response to stimulus, chemical, and stress **(Supplementary Fig.3)**. On the contrary, the bottom 50 affected HVGs by CCC were enriched in immune cell-specific biological processes, such as T cell-mediated cytotoxicity, natural killer cell-mediated cytotoxicity, lymphocyte differentiation, mononuclear cell differentiation, MHC protein complex assembly, and cell development **(Supplementary Fig.3)**. The enrichment results showed that scDCA accurately deciphered the expression profiles of target genes.

**Figure 3:**
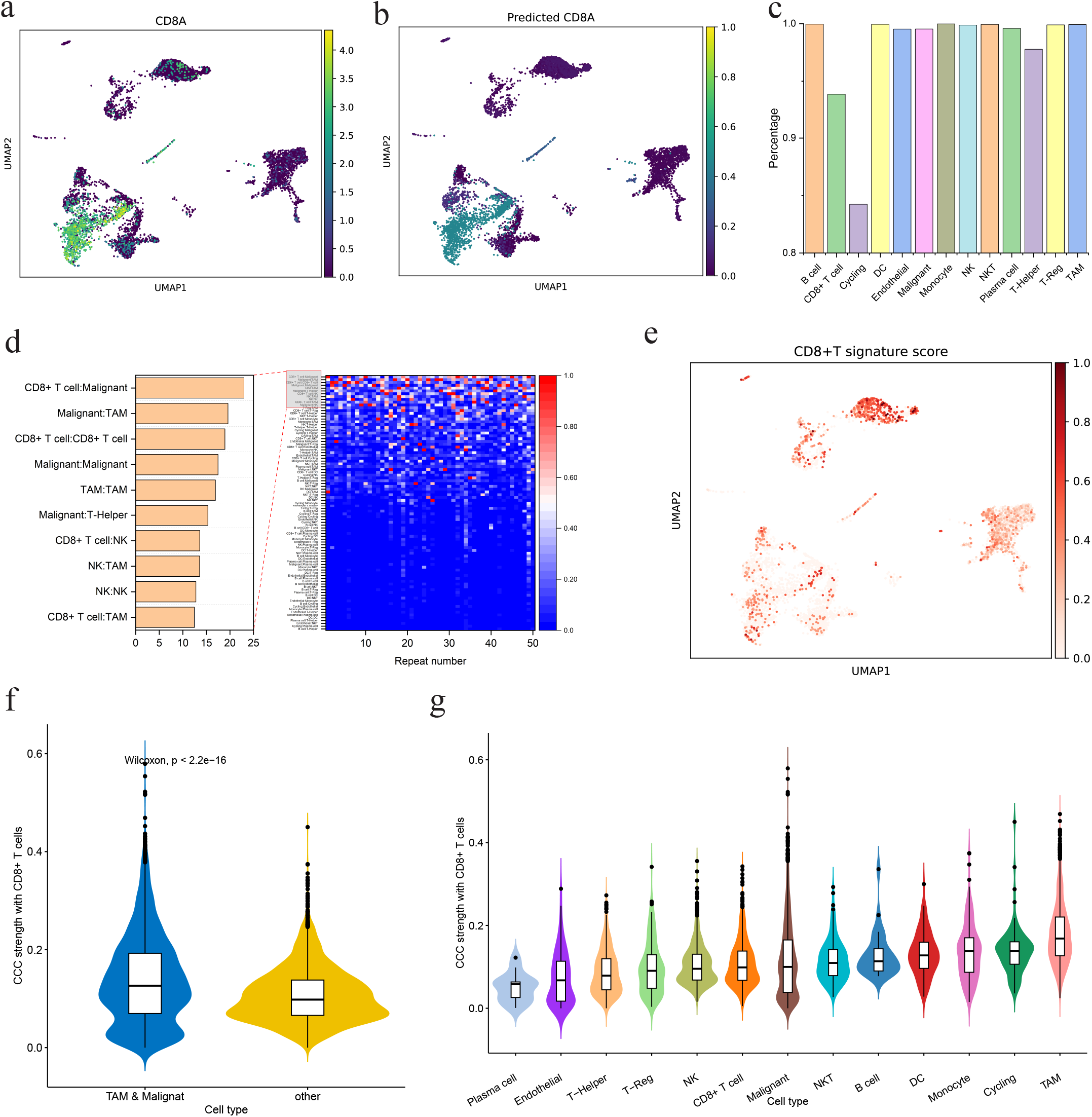
The DCA that regulates the CD8A expression pattern is accurately deciphered. **(a)** UMAP of cell samples, followed by the real expression pattern of the CD8A gene in UMAP space. **(b)** UMAP of cell samples, followed by the predicted expression of CD8A gene by scDCA. **(c)** The ratio of Δ*E* to the total expression levels (Δ*E* and *E*_0_) in each cell type. **(d)** The corresponding cell type pairs of the top 10 view (169 views in total) with the highest attention weights in the model predicting CD8A expression profile. The heatmap displays the attention weights of each view obtained from repeated the training procedure 50 times. The bar plot represents the mean values of the attention weights of each view. **(e)** UMAP of cell samples, followed by the CD8+ T signature score in UMAP space. **(f)** Comparison of communication strength of CD8+ T cells and malignant/TAM cells with that of CD8+ T cells and other cell types. **(g)** Comparison of communication strength with CD8+ T cells among each cell types.

Furthermore, scDCA prioritized the DCA of affecting CD8A expression based on the model’s attention value (see section 2.9 for details), with the CD8+ T cell-Malignant and Malignant-TAM communication ranked higher, suggesting that they dominate in affecting CD8A expression (**Fig. 3.d**). As a mark gene of CD8+ T cells, the DCA affecting CD8A expression also reflects the primary communicating cell types of CD8+ T cells in the TME. On the other hand, prior knowledge has showed that CD70, ICOSLG, CD155 (PVR), CD112 (NECTIN2), and PD-L1 genes are regulatory molecules of T cells [33]. Based on this, we calculated the average expression of them (called CD8+ T signature score) and observed its high expression in malignant cells, which was consistent with the results of scDCA (**Fig. 3.e**). Previous studies have aslo found that malignant cells induced macrophages to express high levels of IL-15R*a*^+^, which reduced the protein levels of chemokine CX3CL1 in malignant cells to inhibit the recruitment of CD8+ T cells [34]. Besides, malignant cells with stem-like states were revealed to drive T cell dysfunction via immunosuppressive macrophages [35]. These results all suggested strong CCCs between CD8+ T and malignant cells, which supported the results of scDCA.

To further compare the communication strength of CD8+ T cells and malignant/TAM cells with that of CD8+ T cells and other cell types in the TME, we calculated the average communication probability with all CD8+ T cells for each single cell. The comparison supported that malignant and TAM cells had significantly higher communication strength with CD8+ T cells than the rest cell types (**Fig. 3.f**). In terms of each cell type, TAM still had the highest communication strength, while malignant cells had a lower median value but a wider range (**Fig. 3.g**). For malignant cells, we further divided them into two groups based on their average communication strength with CD8+ T cells. The group with higher communication strength had accordingly higher activity of antigen presentation via MHC I and IFN-*y* signaling **(Supplementary Fig.4)**. These observations suggested strong cellular communication between a subset of malignant cells and CD8+ T cells. All of the above investigations revealed the dominant CCC between CD8+ T and malignant/TAM cells which again supported the results of scDCA.

### 3.4. The DCA that affects the key regulatory genes in tumor-associated macrophages is accurately deciphered

In section 3.3, we found that TAMs have strong inter-cellular communications with CD8+ T cells, which made us interested in the DCA associated with TAMs. We further deciphered several key regulatory genes in TAMs, including FOLR2 (Folate receptor *β*), APOE (Apolipoprotein E), and FABP5 (Fatty Acid Binding Protein 5) [36, 37, 38]. A multi-view CCC network centered on the TAMs was constructed, resulting in 13 views that involved the communications between other 12 cell types and TAMs as well as TAM themselves. Then, scDCA was retrained and priority sorted on dominant cellular communication based on attention weight in anticipation of revealing holographic networks for specific TAM markers. After training, scDCA similarly accurately reconstructed the gene expression profiles, which were highly consistent with the real gene expression profiles (**Supplementary Fig.5**).

According to the analysed results, the expression of FOLR2 in TAMs was mainly affected by the communication between malignant cells and TAMs, while APOE and FABP5 were greatly affected by the communication within TAM cells (**Fig. 4.a**). FOLR2 was canonically described as a marker of macrophages M2 polarization [36, 39], and indeed specifically over-expressed in part of the TAMs (**Supplementary Fig.5.a**). In the TME, M2 polarization is usually associated with the tumor-promoting phenotype of TAMs [40]. APOE was found to be specifically over-expressed in TAMs in the previous analysis about gastric and breast cancers [37, 41], consistently, we also observed its high expression in TAMs of our ccRCC dataset (**Supplementary Fig.5.b**). It also verified as a potential prognostic biomarker for ccRCC recurrence by [42]. Besides, previous analysis supported that FABP5 controlled macrophage alternative activation, also, FABP5-expressing TAMs were present at invasive cancer regions, produced various immunoregulatory molecules (including PD-L1 and PD-L2) [40, 43].

**Figure 4:**
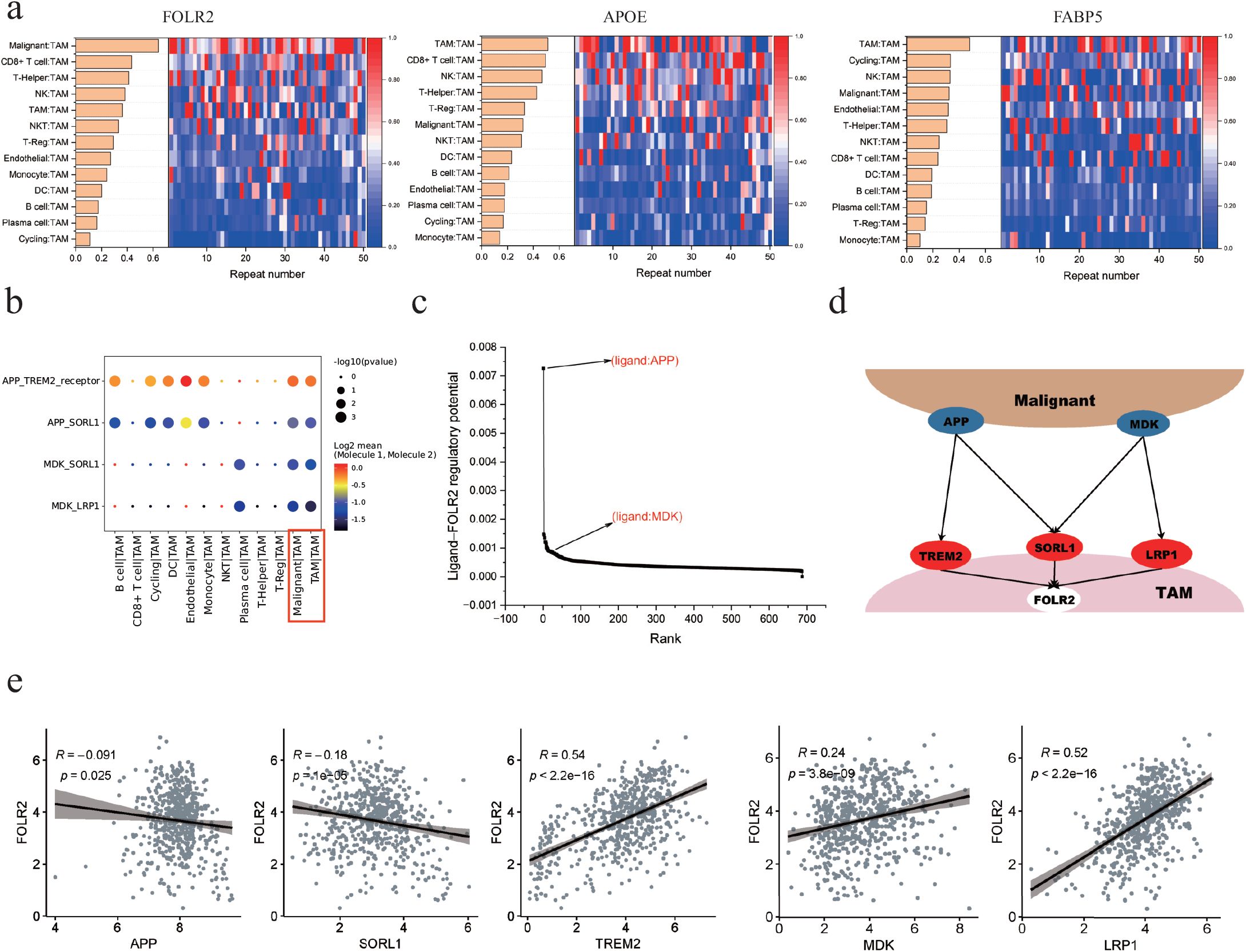
The DCA that affects the key regulatory in tumor-associated macrophages is accurately deciphered. **(a)** Rank of corresponding cell type pairs (views, 13 in total) based on the attention weights in the model predicting expression profile of the key factors FOLR2, APOE, FABP5 in TAMs respectively. The heatmap displayed the attention weights of each view obtained from repeating the training procedure 50 times. The bar plot represents the average attention weights of each view. **(b)** The significant L-R interactions between malignant cells and TAMs which assessed by the CellPhoneDB analysis. **(c)** Rank of ligands based on the regulatory potential on target gene FOLR2 calculated by the priori database NicheNet. **(d)** The schematic diagram of ligands APP, and MDK from malignant cells bind to the receptors TREM2, SORL1, and LRP1 respectively to coordinately regulate the expression of FOLR2 in TAMs. **(e)** The Pearson’s correlation analysis of expression between ligand or receptor genes and FOLR2 in TCGA KIRC cohort.

Further, taking FOLR2 as an example, we further analyzed it from the ligand-receptor-target (L-R-T) perspective. We integrated L-R-T regulatory data from CellPhoneDB and NicheNet to explore important L-R pairs that regulated FOLR2 expression in TAMs and the relationship between these L-R pairs and malignant cells. In the results, Cell-PhoneDB revealed L-R pairs of APP:TREM2, APP:SORL1, MDK:SORL1 and MDK:LRP1 significantly mediated the communication between malignant cells and TAMs as well as TAMs themselves (**Fig. 4.b**). NicheNet evaluated the potential of different ligands to regulate FOLR2. The ligands APP and MDK were at the top of the list, especially APP was the most effective ligand in regulating FOLR2 (**Fig. 4.c**). Specifically, ligands APP and MDK were highly expressed in the malignant cells while the corresponding receptors TREM2, SORL1, and LRP1 were highly expressed in the TAMs (**Supplementary Fig.6**), therefore, they might jointly mediate the regulation of FOLR2 expression in TAMs (**Fig. 4.d**. In addition, co-expression analysis in the bulk RNA-seq samples from TCGA also supported the regulation of these L-R pairs on FOLR2 (**Fig. 4.e**). It is worth noting that APP and MDK regulate different receptors in different directions, which in turn results in a superposition of multiple effects on FOLR2. All the results demonstrate the accuracy and reliability of scDCA for deciphering the DCA that regulates specific target genes and facilitates further effective excavation of mediating L-R pairs.

### 3.5. The DCA that affects functional states of malignant cells is accurately deciphered

Compelling evidence highlighted that multiple CCCs between different cell types were involved in affecting functional states of cells, such as angiogenesis [44], EMT [45], and invasion [46] of malignant cells. scDCA could be also used to decipher the impact of each communicating cell type pairs on the cell states. To do this, we first calculated the activity scores for 14 functional states of 7393 malignant cells in our P76_scRNA data. There was a high degree of functional heterogeneity among these malignant cells, with a small fraction of them exhibiting elevated levels of epithelial-mesenchymal transition (EMT), invasion, metastasis, and hypoxia, suggesting a higher malignancy potential in this subpopulation (**Supplementary Fig.7**).

Then, scDCA was retrained based on the reconstructed multi-view CCC network centered on the malignant cells, resulting in 13 views that involved the communications between 12 other cell types and malignant cells as well as malignant cells themselves. Different from the gene expression analysis in the previous section, our goal is to predict the functional state scores of malignant cells and assign initial random feature vectors to each individual malignant cell due to the fact that the functional state is associated with the individual cell (**see section 2.7 for details**). In the results, scDCA had significant accuracy in predicting various cell functional states, such as the DNA repair state with a Pearson correlation of 0.875 and p-value *<* 0.05, and the apoptosis state with a Pearson correlation of 0.879 and p-value *<* 0.05 (see **Supplementary Table.1** for more details).

Similarly based on the average attention weight of 50 repetitions of training, scDCA deciphered the DCA that affected the functional state of malignant cells as shown in **Fig. 5.a**. Overall, natural killer (NK), T-helper, malignant itself, CD8+ T, TAMs, endothelial and monocyte cells had a greater impact on the functional states of malignant cells. In contrast, B and plasma cells had less overall effect. From the results, the cell states that were significantly influenced by the malignant cells themselves were angiogenesis and differentiation. Indeed, we observed VEGFA (vascular endothelial growth factor), a major signal that influences angiogenesis and promotes differentiation of malignant cells [47, 48], is highly specific expression in malignant cells **(Fig. 5.b)**. In addition, NK cells affected various functional states of malignant cells, especially their cell cycle and differentiation. This phenomenon has been shown by recent studies in which NK cells upregulated the expression of specific genes to induce cell cycle or differentiation of malignant cells [49, 50, 51], which further corroborates the results of scDCA.

**Figure 5:**
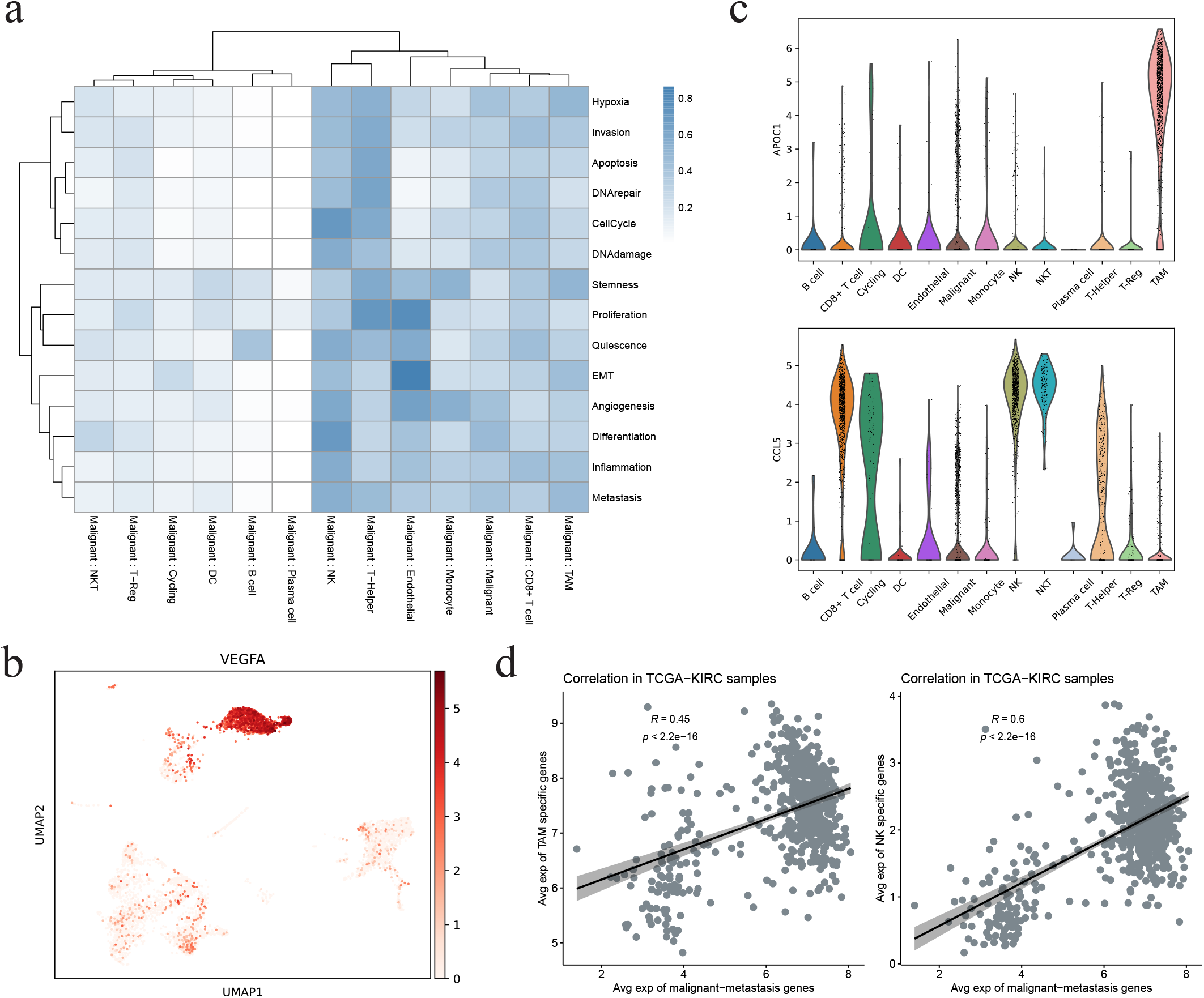
The DCA that affects functional states of malignant cells is accurately deciphered. **(a)** Rank of attention weights of corresponding cell type pairs (views) between malignant cells and other cell types in the model predicting the functional states of malignant cells. The heatmap displayed the attention weights of each view obtained from repeated the training procedure 50 times. The bar plot represented the average attention weights of each view. **(b)** UMAP of cell samples, followed by the expression pattern of VEGFA. **(c)** Expression distribution of the two metastatic regulators (TAM-specific gene APOC1 and NK-specific gene CCL5) in different cell types. **(d)** Correlation analysis between signature of “invasion-metastasis cascade” states of malignant cells and marker genes of TAMs and NK cells in the TCGA-KIRC samples (N=607).

Furthermore, tumor invasion and metastasis involve a sophisticated cascade process, often referred to as the “invasion-metastasis cascade”, and EMT has been shown to be a critical first step for the cascade process [52, 53, 54]. In our results, the CCC between malignant and endothelial cells was the most influential factor for the EMT state of malignant cells, this finding was strongly supported by the prior knowledge about the EMT process [55]. Considering the three cascade states comprehensively, we observed that endothelials, NK, T-helper, and TAM cells were the dominant cell types that were involved in regulating them (**Supplementary Fig.8**). Intersection of the dominant cell types showed NK cells and TAMs might exert common effects for the “invasion-metastasis cascade” of malignant cells, which had been validated by substantial studies. NK cells play a critical role in the control of metastasis of malignant cells as they mediate metastatic cell immunoediting, [56, 57], and IFN-*γ*-dependent NK cell activation is vital to suppress malignant metastasis [58]. Also, perturbation of the EMT state of malignant cell lines altered several representative NK cell-regulatory ligands [59]. Similarly, TAM cells was found to promote malignant progression by enhancing invasion and metastasis [60, 61, 62]. Specifically, M2-polarized TAM was found to promote EMT and activate invasion programs in malignant cells [63]. The expression pattern of some specific metastatic regulators also supported the critical influence of NK and TAMs cells on the “invasion-metastasis cascade” states of malignant cells (**Fig. 5.c**). In particular, Apolipoprotein C-I (APOC1) has been identified as a novel pro-metastatic factor [64], which enhances the metastasis of malignant cells via EMT pathway in ccRCC and gastric cancer [65, 66]. Indeed, we observed that APOC1 was specifically highly expressed on TAMs in our data, which supports the potential of TAMs in promoting malignant cell metastasis as revealed by scDCA (**Fig. 5.c**). Another example is the NK-derived chemokine CCL5, which promotes malignant metastasis and has been verified as an important mediator for immune evasion in circulating malignant cells [67, 68, 69]. Finally, we analyzed the correlation between “invasion-metastasis cascade” signature genes and TAM/NK cell marker genes at the bulk level. We first filtered the three sets of “invasion-metastasis cascade” associated genes based on the malignant cell-specific expressed genes. Their average expression was found significantly correlated in TCGA-KIRC samples (N=607) (**Fig. 5.d**). These collective findings provide compelling evidence that scDCA possesses the capability to decipher intercellular communication that affects the functional status of malignant cells in the TME.

### 3.6. The DCA associated with clinical response of immune checkpoint blockade is accurately deciphered

Understanding the alteration of DCA under therapeutic interventions is crucial for dissecting the specific molecular mechanism of disease progression or therapeutic response. To compare with the previously analyzed untreated patient P76, we obtained scRNA-seq data from another patient with advanced ccRCC, P915, who received ICB therapy and exhibited a clinical response [24] (**Supplementary Table 2**). Totally 6304 cells were obtained and 14 cell types were annotated according to the original publication **(Fig. 6.a)**. We can see that there was a significant increase of CD8+ T cells after ICB treatment **(Fig. 6.b)**.

**Figure 6:**
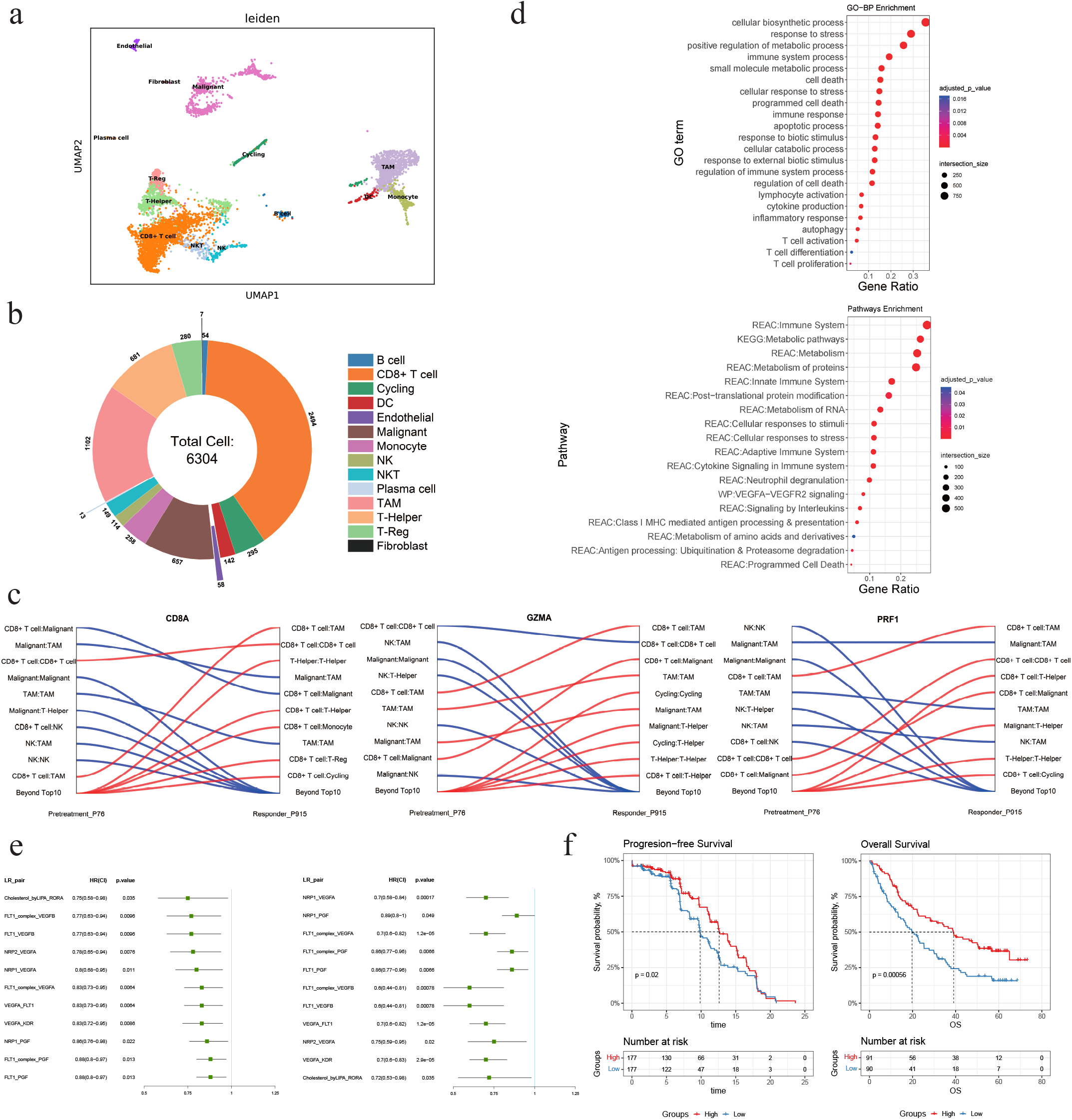
The DCA associated with clinical response of immune checkpoint blockade is accurately deciphered. **(a)** UMAP of all captured cell samples from patient P915. Cell type annotation was inherited from the data resource analysis and visualized with different colors. **(b)** The proportions of all cell types in the P915_scRNA dataset. **(c)** Mapping of the rank changes of DCA for 3 signature genes from comparing ICB-responder (P915) to untreated patients (P76). **(d)** Functional enrichment analysis of GO-BP (up) and KEGG pathways (down) for the differential expressed genes of TAMs derived from comparing the ICB-responder (P915) to the untreated patient (P76). **(e)** Forest plot of the L-R pairs that exhibited significant prognostic significance (univariate Cox regression analysis) in two independent advanced ccRCC cohorts following ICB immunotherapy. Left: data from ICB immunotherapy-only subpopulations of Motzer_NatMed_2020 cohort; Right: data from ICB immunotherapy-only subpopulations of Braun_NatMed_2020_Checkmate025 cohort. **(f)** Kaplan-Meier curves and log-rank test for the integration of the 11 L-R pairs in the two ICB-treatment-only subpopulations. The average expression of these LR pairs was summed up for the prognostic association analysis. Left: data from ICB immunotherapy-only subpopulations of Motzer_NatMed_2020 cohort; Right: data from ICB immunotherapy-only subpopulations of Braun_NatMed_2020_Checkmate025 cohort.

The scDCA was then applied to decipher the DCA that affects CD8A expression pattern in ICB-responder P915 scRNA-seq data. Compared to the untreated patient (**Fig. 3.d**), the top DCA changed from CD8+ T cell-malignant to CD8+ T cell-TAM **(Fig. 6.c)**. To further investigate the DCA alteration in the ICI-responders, we analyzed two additional genes, GZMA and PRF1, which are signature genes of cytolytic activity, indicating the ability of CD8+ T cells to clear malignant cells [70, 71]. Notably, similar to CD8A, DCA that affects these two cytolytic indicators also turned into CD8+ T cell-TAM **(Fig. 6.c)**. These findings were supported by the previous observation [24] that normalized effect or molecule (e.g., IFNG) levels were robustly correlated with TAM fractions and expression signatures derived from comparing ICB (immune checkpoint blockade)-exposed to ICB-naive TAMs. All these results suggested the increased potential of TAM in regulating CD8+ T cells in ICI-responders compared to untreated patient.

Additionally, the scDCA was also applied to decipher the DCA that affected 3 cellular signatures associated with the clinical response of ICB [24], including the tumor program 1 (TP1) and tumor program 2 (TP2) active in malignant cells that may drive interactions with the immune system, as well as the signature of immune checkpoint and evasion (**Supplementary Fig.9**). We found that the TAM-malignant cell pair significantly rose to the top 3 ranks related to treatment response in ICB-responders compared to untreated patient, suggesting that TAMs affected the therapeutic responsiveness of malignant cells.

Therefore, we speculated that the CCC changes generated by TAM after ICB therapeutic intervention are crucial in mediating immunotherapy response. On the basis, we then obtained the differential expressed genes of TAMs derived from comparing ICB-responder (P915) to the untreated patient (P76). Functional enrichment analysis of GO-biological process revealed the up-regulated genes after ICB treatment were related to several processes involving immune response and cell death, such as “programmed cell death”, “apoptotic process”, “autophagy”, “inflammatory response”, “proliferation, differentiation and activation of T cell” and “cytokine production” **(Fig. 6.d)**. In terms of pathways analysis, similar to GO-BP results, the immune system, antigen processing and presentation, cytokine signaling, and programmed cell death were significantly enriched. Besides, metabolic pathways and VEGFA:VEGFR2 signaling were also enriched **(Fig. 6.d)**. These results indicated the reinforced immune regulating potential of TAM cells in ICB-responder dataset.

Furthermore, the differences in L-R interactions between TAMs and CD8+ T cells were further compared. To our surprise, there was a significant increase in L-R pairs in the ICB-responder dataset compared to the untreated one. In specific, according to cellPhoneDB analysis, there were 62 L-R pairs between TAMs and CD8+ T cells in the untreated patient while 1236 L-R pairs in the ICB-responder patient, respectively. For the 1028 different LR pairs, we further investigated their correlation with the clinical benefit of the ICI treatment. Using the average expression of each LR pair, we found that 11 LR pairs exhibited significant prognostic significance in two independent advanced ccRCC cohorts following ICB immunotherapy **(Fig. 6.e)**. Importantly, their prognostic association was significant only within subpopulations receiving ICB immunotherapy, compared to Sunitinib [31] and mTOR inhibitor everolimus [32] **(Supplementary Fig.10.a and b)**. Furthermore, when combining the average expression of these 11 L-R pairs, a similar prognostic association was observed in ICIB-treatment-only subpopulations but not the targeted therapy subpopulations **(Fig. 6.f, Supplementary Fig.10.c and d)**. In summary, we attempted to apply scDCA in patients with different clinical phenotypes and explored the alteration in CCC that have a unique contribution to clinical intervention and patient prognosis. In the comparison of untreated and ICI-responder patients of advanced RCC, scDCA helped discern the prominent role of TAM in regulating the response of CD8+ T cells after immunotherapy and the emerged L-R pairs showed potential that mediating clinical benefit of immunotherapy.

## 4. Discussion and Conclusions

In response to the computational challenge of deciphering the downstream functional impact of cell-cell communication, we proposed a computational method, scDCA. It can innovatively prioritize cell type pairs that play a dominant role in downstream functional impacts (called DCA). In particular, scDCA builds the comprehensive structure of CCC at the single cell resolution to maximally leverage the information from communicative heterogeneity and specificity. We construct the multi-view graph convolution network based on the scRNA-seq data to model the impact of different cell type pairs on the target gene expression or functional states of cells. In advanced renal cell carcinoma dataset, scDCA reveals the DCA that affects the key factors in CD8+ T cells and TAMs as well as the multiple functional states of malignant cells. In addition, it also provides insights into changes in DCA before and after the immunotherapy so as to facilitate understanding of the intercellular communication basis of the clinical response.

Current existing methods such as CellChat, CellPhoneDB, etc., perform CCC analysis at the level of the cell type or cluster, discarding single-cell-level information. However, biologically, CCC does not act at the group level, but between individual cells, and not all cells have a direct or equal impact on downstream functional effect. A case study of scRNA-seq data from a ccRCC patient indicated that the communication strength, the count of communication types, and the ligand-receptor pair type with maximum communication strength varied at the single cell resolution. Hence, to leverage the full information contained within scRNA-seq data by looking at the communication landscape between individual cells, scDCA utilized graph model to model the interaction between each cell in a pair of cell types. Graph neural network (GNN) has excellent performance on graph-structured data. Multi-view GNN framework is applicable to model the holographic network of CCC, and attention mechanisms facilitate biological interpretability of which cell type pairs play the major role in the specific functional process. A case study of specific CD8A expression patterns in the P76_scRNA data indicated that scDCA can accurately reconstruct gene expression, decipher the dominant cell communication assembly affecting specific gene expression, and the ratio of the expression change caused by CCC in each cell type. A case study of the key factors in TAMs indicated that scDCA can also effectively decipher intercellular communications between specific cell types. Furthermore, a case study of functional states of malignant cells in the P76_scRNA data indicated that scDCA possesses the capability to elucidate intercellular communication that influences the functional status of specific cell types.

Longitudinal datasets pose an additional opportunity and challenge for comparative analyses because there is a priori knowledge about the opposite clinical phenotype between different samples. While previous analyses have mostly integrated single-cell samples from patient cohorts and observed communication between cell types in the integrated patient collection, scDCA allows for a more fine-grained and targeted characterization of intercellular communication from the perspective of a single patient (patient-individual-specific characterization of intercellular communication effects). Moreover, single-cell samples from multiple patient sources offer the possibility to compare cell communication-specific changes under different clinical phenotypes or therapeutic intervention. In a case study of the clinical response of immune checkpoint blockade, we compared differences in intercellular communication between patients with different clinical phenotypes and dissected changes in cellular communication effects that impact the regulation of specific key functional genes and pathways.

In future work, we intend to incorporate spatial information into our method with the development of spatial sequencing technologies. Hence, more subdivided cell types and positions on pseudotime trajectories can be utilized to obtain a much more comprehensive cellular landscape. Furthermore, prior knowledge is also considered in the future, such as the intracellular signaling networks or known biological functions of some ligands, to better decipher intercellular communication and gain a deeper biological understanding.

## Supporting information

Supplemental results

## 5. Data Availability

- This paper analyzes existing, publicly available data. The details of the datasets are displayed in the section 2.1.
- All original code and the preprocessed data have been deposited at GitHub (https://github.com/pengsl-lab/scDCA.git) and is publicly available as of the date of publication. Concurrently, we have added the description of how to use our program.
- Any additional information required to reanalyze the data reported in this paper is available from the lead contact upon request.

## 7. Declaration of Interests

The authors declare that they have no competing interests.

## 8. Author Contributions

All authors were involved in the conceptualization of the scDCA method. LWX and SLP conceived and supervised the project. BYJ and LWX designed the study and developed the approach. XQW and XW collected the data. BYJ and LWX analyzed the results. BYJ, XQW, XW, LWX and SLP contributed to the review of the manuscript before submission for publication. All authors read and approved the final manuscript.

## 9. Funding

NSFC-FDCT Grants 62361166662; National Key R&D Program of China 2023YFC3503400, 2022YFC3400400; Key R&D Program of Hunan Province 2023GK2004, 2023SK2059, 2023SK2060; Top 10 Technical Key Project in Hunan Province 2023GK1010; Key Technologies R&D Program of Guangdong Province (2023B1111030004 to FFH). The Funds of State Key Laboratory of Chemo/Biosensing and Chemometrics, the National Supercomputing Center in Changsha (http://nscc.hnu.edu.cn/), and Peng Cheng Lab.

## 10. Acknowledgements

We thank professor Fengxiang Gao, Li Zhang and Ethan Williams for constructive comments of this work. We thank Dr. Xiongjun Zhao for suggestions on algorithm improvement and preprocessing of scRNA-seq data.

## Notes

### Competing Interest Statement

The authors have declared no competing interest.

## References

[1] E. Armingol, A. Officer, O. Harismendy, N. E. Lewis, Deciphering cell–cell interactions and communication from gene expression, Nature Reviews Genetics 22 (2) (2021) 71–88.

[2] H. Zhang, X. Lin, Z. Han, L.-J. Qu, J. Chai, Crystal structure of pxytdif complex reveals a conserved recognition mechanism among cle peptide-receptor pairs, Cell research 26 (5) (2016) 543–555.

[3] Y. He, R. M. Rodrigues, X. Wang, W. Seo, J. Ma, S. Hwang, Y. Fu, E. Trojnár, C. Mátyás, S. Zhao, et al., Neutrophil-to-hepatocyte communication via ldlr-dependent mir-223–enriched extracellular vesicle transfer ameliorates nonalcoholic steatohepatitis, The Journal of clinical investigation 131 (3) (2021).

[4] Y. Suhail, M. P. Cain, K. Vanaja, P. A. Kurywchak, A. Levchenko, R. Kalluri, et al., Systems biology of cancer metastasis, Cell systems 9 (2) (2019) 109–127.

[5] A. A. Almet, Z. Cang, S. Jin, Q. Nie, The landscape of cell–cell communication through single-cell transcriptomics, Current opinion in systems biology 26 (2021) 12–23.

[6] D. S. Chen, I. Mellman, Elements of cancer immunity and the cancerimmune set point, Nature 541 (7637) (2017) 321–330.

[7] R. Kalluri, The biology and function of fibroblasts in cancer, Nature Reviews Cancer 16 (9) (2016) 582–598.

[8] Y. Yang, G. Li, Y. Zhong, Q. Xu, Y.-T. Lin, C. Roman-Vicharra, R. S. Chapkin, J. J. Cai, sctenifoldxct: A semi-supervised method for predicting cell-cell interactions and mapping cellular communication graphs, Cell Systems 14 (4) (2023) 302–311.

[9] M. P. Kumar, J. Du, G. Lagoudas, Y. Jiao, A. Sawyer, D. C. Drummond, D. A. Lauffenburger, A. Raue, Analysis of single-cell rna-seq identifies cell-cell communication associated with tumor characteristics, Cell reports 25 (6) (2018) 1458–1468.

[10] D. Dimitrov, D. Türei, M. Garrido-Rodriguez, P. L. Burmedi, J. S. Nagai, C. Boys, R. O. Ramirez Flores, H. Kim, B. Szalai, I. G. Costa, et al., Comparison of methods and resources for cell-cell communication inference from single-cell rna-seq data, Nature communications 13 (1) (2022) 3224.

[11] M. Efremova, M. Vento-Tormo, S. A. Teichmann, R. Vento-Tormo, Cellphonedb: inferring cell–cell communication from combined expression of multi-subunit ligand–receptor complexes, Nature protocols 15 (4) (2020) 1484–1506.

[12] S. Jin, C. F. Guerrero-Juarez, L. Zhang, I. Chang, R. Ramos, C.-H. Kuan, P. Myung, M. V. Plikus, Q. Nie, Inference and analysis of cellcell communication using cellchat, Nature communications 12 (1) (2021) 1088.

[13] R. Browaeys, W. Saelens, Y. Saeys, Nichenet: modeling intercellular communication by linking ligands to target genes, Nature methods 17 (2) (2020) 159–162.

[14] Y. Hu, T. Peng, L. Gao, K. Tan, Cytotalk: De novo construction of signal transduction networks using single-cell transcriptomic data, Science Advances 7 (16) (2021) eabf1356.

[15] R. Hou, E. Denisenko, H. T. Ong, J. A. Ramilowski, A. R. Forrest, Predicting cell-to-cell communication networks using natmi, Nature communications 11 (1) (2020) 5011.

[16] D. Pham, X. Tan, J. Xu, L. F. Grice, P. Y. Lam, A. Raghubar, J. Vukovic, M. J. Ruitenberg, Q. Nguyen, stlearn: integrating spatial location, tissue morphology and gene expression to find cell types, cell-cell interactions and spatial trajectories within undissociated tissues, BioRxiv (2020) 2020–05.

[17] L. Garcia-Alonso, L.-F. Handfield, K. Roberts, K. Nikolakopoulou, R. C. Fernando, L. Gardner, B. Woodhams, A. Arutyunyan, K. Polanski, R. Hoo, et al., Mapping the temporal and spatial dynamics of the human endometrium in vivo and in vitro, Nature Genetics 53 (12) (2021) 1698–1711.

[18] A. J. Wilk, A. K. Shalek, S. Holmes, C. A. Blish, Comparative analysis of cell–cell communication at single-cell resolution, Nature Biotechnology (2023) 1–14.

[19] D. S. Fischer, A. C. Schaar, F. J. Theis, Modeling intercellular communication in tissues using spatial graphs of cells, Nature Biotechnology 41 (3) (2023) 332–336.

[20] Y. Yuan, Z. Bar-Joseph, Gcng: graph convolutional networks for inferring gene interaction from spatial transcriptomics data, Genome biology 21 (1) (2020) 1–16.

[21] A. Vaswani, N. Shazeer, N. Parmar, J. Uszkoreit, L. Jones, A. N. Gomez, Ł. Kaiser, I. Polosukhin, Attention is all you need, Advances in neural information processing systems 30 (2017).

[22] G. Chen, Z.-P. Liu, Graph attention network for link prediction of gene regulations from single-cell rna-sequencing data, Bioinformatics 38 (19) (2022) 4522–4529.

[23] H. Li, T. Ma, M. Hao, W. Guo, J. Gu, X. Zhang, L. Wei, Decoding functional cell–cell communication events by multi-view graph learning on spatial transcriptomics, Briefings in Bioinformatics 24 (6) (2023) bbad359.

[24] K. Bi, M. X. He, Z. Bakouny, A. Kanodia, S. Napolitano, J. Wu, G. Grimaldi, D. A. Braun, M. S. Cuoco, A. Mayorga, et al., Tumor and immune reprogramming during immunotherapy in advanced renal cell carcinoma, Cancer Cell 39 (5) (2021) 649–661.

[25] Z. Liu, D. Sun, C. Wang, Evaluation of cell-cell interaction methods by integrating single-cell rna sequencing data with spatial information, Genome Biology 23 (1) (2022) 1–38.

[26] F. Noël, L. Massenet-Regad, I. Carmi-Levy, A. Cappuccio, M. Grand-claudon, C. Trichot, Y. Kieffer, F. Mechta-Grigoriou, V. Soumelis, Dissection of intercellular communication using the transcriptome-based framework icellnet, Nature communications 12 (1) (2021) 1089.

[27] R. W. Balluffi, S. M. Allen, W. C. Carter, Kinetics of materials, John Wiley & Sons, 2005.

[28] F. A. Wolf, P. Angerer, F. J. Theis, Scanpy: large-scale single-cell gene expression data analysis, Genome biology 19 (2018) 1–5.

[29] T. N. Kipf, M. Welling, Semi-supervised classification with graph convolutional networks, arXiv preprint 1609.02907 (2016).

[30] H. Yuan, M. Yan, G. Zhang, W. Liu, C. Deng, G. Liao, L. Xu, T. Luo, H. Yan, Z. Long, et al., Cancersea: a cancer single-cell state atlas, Nucleic acids research 47 (D1) (2019) D900–D908.

[31] R. J. Motzer, P. B. Robbins, T. Powles, L. Albiges, J. B. Haanen, J. Larkin, X. J. Mu, K. A. Ching, M. Uemura, S. K. Pal, et al., Avelumab plus axitinib versus sunitinib in advanced renal cell carcinoma: biomarker analysis of the phase 3 javelin renal 101 trial, Nature medicine 26 (11) (2020) 1733–1741.

[32] D. A. Braun, Y. Hou, Z. Bakouny, M. Ficial, M. Sant’Angelo, J. Forman, P. Ross-Macdonald, A. C. Berger, O. A. Jegede, L. Elagina, et al., Interplay of somatic alterations and immune infiltration modulates response to pd-1 blockade in advanced clear cell renal cell carcinoma, Nature medicine 26 (6) (2020) 909–918.

[33] S. Jeong, S.-H. Park, Co-stimulatory receptors in cancers and their implications for cancer immunotherapy, Immune Network 20 (1) (2020).

[34] W. Zhang, Q. Zhang, N. Yang, Q. Shi, H. Su, T. Lin, Z. He, W. Wang, H. Guo, P. Shen, Crosstalk between il-15ra+ tumorassociated macrophages and breast cancer cells reduces cd8+ t cell recruitment, Cancer Communications 42 (6) (2022) 536–557.

[35] Y. Tang, D. J. Kwiatkowski, E. P. Henske, Midkine expression by stem-like tumor cells drives persistence to mtor inhibition and an immune-suppressive microenvironment, Nature Communications 13 (1) (2022) 5018.

[36] A. Puig-Kröger, E. Sierra-Filardi, A. Domínguez-Soto, R. Samaniego, M. T. Corcuera, F. Gómez-Aguado, M. Ratnam, P. Sánchez-Mateos, A. L. Corbí, Folate receptor β is expressed by tumor-associated macrophages and constitutes a marker for m2 anti-inflammatory/regulatory macrophages, Cancer research 69 (24) (2009) 9395–9403.

[37] P. Zheng, Q. Luo, W. Wang, J. Li, T. Wang, P. Wang, L. Chen, P. Zhang, H. Chen, Y. Liu, et al., Tumor-associated macrophages-derived exosomes promote the migration of gastric cancer cells by transfer of functional apolipoprotein e, Cell death & disease 9 (4) (2018) 434.

[38] J. Hao, F. Yan, Y. Zhang, A. Triplett, Y. Zhang, D. A. Schultz, Y. Sun, J. Zeng, K. A. Silverstein, Q. Zheng, et al., Expression of adipocyte/macrophage fatty acid–binding protein in tumor-associated macrophages promotes breast cancer progression, Cancer research 78 (9) (2018) 2343–2355.

[39] E. Sierra-Filardi, A. Puig-Kröger, F. J. Blanco, C. Nieto, R. Bragado, M. I. Palomero, C. Bernabéu, M. A. Vega, A. L. Corbí, Activin a skews macrophage polarization by promoting a proinflammatory phenotype and inhibiting the acquisition of anti-inflammatory macrophage markers, Blood, The Journal of the American Society of Hematology 117 (19) (2011) 5092–5101.

[40] M. J. Pittet, O. Michielin, D. Migliorini, Clinical relevance of tumour-associated macrophages, Nature reviews Clinical oncology 19 (6) (2022) 402–421.

[41] R. N. Ramos, Y. Missolo-Koussou, Y. Gerber-Ferder, C. P. Bromley, M. Bugatti, N. G. Núñez, J. T. Boari, W. Richer, L. Menger, J. Denizeau, et al., Tissue-resident folr2+ macrophages associate with cd8+ t cell infiltration in human breast cancer, Cell 185 (7) (2022) 1189–1207.

[42] A. Obradovic, N. Chowdhury, S. M. Haake, C. Ager, V. Wang, L. Vlahos, X. V. Guo, D. H. Aggen, W. K. Rathmell, E. Jonasch, et al., Single-cell protein activity analysis identifies recurrence-associated renal tumor macrophages, Cell 184 (11) (2021) 2988–3005.

[43] Y. Hou, D. Wei, Z. Zhang, H. Guo, S. Li, J. Zhang, P. Zhang, L. Zhang, Y. Zhao, Fabp5 controls macrophage alternative activation and allergic asthma by selectively programming long-chain unsaturated fatty acid metabolism, Cell Reports 41 (7) (2022).

[44] B. M. Lopes-Bastos, W. G. Jiang, J. Cai, Tumour–endothelial cell communications: important and indispensable mediators of tumour angiogenesis, Anticancer research 36 (3) (2016) 1119–1126.

[45] C. E. Hills, E. Siamantouras, S. Smith, P. Cockwell, K.-K. Liu, P. E. Squires, Tgfβ modulates cell-to-cell communication in early epithelial-to-mesenchymal transition, Diabetologia 55 (2012) 812–824.

[46] F. Calvo, E. Sahai, Cell communication networks in cancer invasion, Current opinion in cell biology 23 (5) (2011) 621–629.

[47] L. Claesson-Welsh, M. Welsh, Vegfa and tumour angiogenesis, Journal of internal medicine 273 (2) (2013) 114–127.

[48] P. Nowak-Sliwinska, J. R. van Beijnum, C. J. Griffioen, Z. R. Huinen, N. G. Sopesens, R. Schulz, S. V. Jenkins, R. P. Dings, F. H. Groenendijk, E. J. Huijbers, et al., Proinflammatory activity of vegf-targeted treatment through reversal of tumor endothelial cell anergy, Angiogenesis 26 (2) (2023) 279–293.

[49] N. K. Wolf, D. U. Kissiov, D. H. Raulet, Roles of natural killer cells in immunity to cancer, and applications to immunotherapy, Nature Reviews Immunology 23 (2) (2023) 90–105.

[50] T. Bald, M. F. Krummel, M. J. Smyth, K. C. Barry, The nk cell–cancer cycle: advances and new challenges in nk cell–based immunotherapies, Nature immunology 21 (8) (2020) 835–847.

[51] A. M. Abel, C. Yang, M. S. Thakar, S. Malarkannan, Natural killer cells: development, maturation, and clinical utilization, Frontiers in immunology 9 (2018) 1869.

[52] A. W. Lambert, D. R. Pattabiraman, R. A. Weinberg, Emerging biological principles of metastasis, Cell 168 (4) (2017) 670–691.

[53] Q. Wei, Y. Qian, J. Yu, C. C. Wong, Metabolic rewiring in the promotion of cancer metastasis: mechanisms and therapeutic implications, Oncogene 39 (39) (2020) 6139–6156.

[54] H. Tian, R. Lian, Y. Li, C. Liu, S. Liang, W. Li, T. Tao, X. Wu, Y. Ye, X. Yang, et al., Akt-induced lncrna val promotes emt-independent metastasis through diminishing trim16-dependent vimentin degradation, Nature communications 11 (1) (2020) 5127.

[55] N. Maishi, K. Hida, Tumor endothelial cells accelerate tumor metastasis, Cancer science 108 (10) (2017) 1921–1926.

[56] K. Nakamura, M. J. Smyth, Immunoediting of cancer metastasis by nk cells, Nature cancer 1 (7) (2020) 670–671.

[57] X. Liu, J. Song, H. Zhang, X. Liu, F. Zuo, Y. Zhao, Y. Zhao, X. Yin, X. Guo, X. Wu, et al., Immune checkpoint hla-e: Cd94-nkg2a mediates evasion of circulating tumor cells from nk cell surveillance, Cancer Cell 41 (2) (2023) 272–287.

[58] Q. Lin, L. Rong, X. Jia, R. Li, B. Yu, J. Hu, X. Luo, S. Badea, C. Xu, G. Fu, et al., Ifn-y-dependent nk cell activation is essential to metastasis suppression by engineered salmonella, Nature Communications 12 (1) (2021) 2537.

[59] H. C. Lo, Z. Xu, I. S. Kim, B. Pingel, S. Aguirre, S. Kodali, J. Liu, W. Zhang, A. M. Muscarella, S. M. Hein, et al., Resistance to natural killer cell immunosurveillance confers a selective advantage to polyclonal metastasis, Nature Cancer 1 (7) (2020) 709–722.

[60] S. Dallavalasa, N. M. Beeraka, C. G. Basavaraju, S. V. Tulimilli, S. P. Sadhu, K. Rajesh, G. Aliev, S. V. Madhunapantula, The role of tumor associated macrophages (tams) in cancer progression, chemoresistance, angiogenesis and metastasis-current status, Current Medicinal Chemistry 28 (39) (2021) 8203–8236.

[61] Y. Chen, Y. Song, W. Du, L. Gong, H. Chang, Z. Zou, Tumor-associated macrophages: an accomplice in solid tumor progression, Journal of biomedical science 26 (1) (2019) 1–13.

[62] A.-M. Georgoudaki, K. E. Prokopec, V. F. Boura, E. Hellqvist, S. Sohn, J. Östling, R. Dahan, R. A. Harris, M. Rantalainen, D. Klevebring, et al., Reprogramming tumor-associated macrophages by antibody targeting inhibits cancer progression and metastasis, Cell reports 15 (9) (2016) 2000–2011.

[63] L. Borriello, A. Coste, B. Traub, V. P. Sharma, G. S. Karagiannis, Y. Lin, Y. Wang, X. Ye, C. L. Duran, X. Chen, et al., Primary tumor associated macrophages activate programs of invasion and dormancy in disseminating tumor cells, Nature communications 13 (1) (2022) 626.

[64] L. Ren, J. Yi, Y. Yang, W. Li, X. Zheng, J. Liu, S. Li, H. Yang, Y. Zhang, B. Ge, et al., Systematic pan-cancer analysis identifies apoc1 as an immunological biomarker which regulates macrophage polarization and promotes tumor metastasis, Pharmacological Research 183 (2022) 106376.

[65] L. An, Y. Liu, Znf460 mediates epithelial-mesenchymal transition to promote gastric cancer progression by transactivating apoc1 expression, Experimental Cell Research 422 (2) (2023) 113452.

[66] Y.-l. Li, L.-w. Wu, L.-h. Zeng, Z.-y. Zhang, W. Wang, C. Zhang, N.-m. Lin, Apoc1 promotes the metastasis of clear cell renal cell carcinoma via activation of stat3, Oncogene 39 (39) (2020) 6203–6217.

[67] Y.-F. Sun, L. Wu, S.-P. Liu, M.-M. Jiang, B. Hu, K.-Q. Zhou, W. Guo, Y. Xu, Y. Zhong, X.-R. Zhou, et al., Dissecting spatial heterogeneity and the immune-evasion mechanism of ctcs by single-cell rna-seq in hepatocellular carcinoma, Nature Communications 12 (1) (2021) 4091.

[68] D. Aldinucci, C. Borghese, N. Casagrande, The ccl5/ccr5 axis in cancer progression, Cancers 12 (7) (2020) 1765.

[69] J. Qiu, L. Xu, X. Zeng, H. Wu, F. Liang, Q. Lv, Z. Du, Ccl5 mediates breast cancer metastasis and prognosis through ccr5/treg cells, Frontiers in Oncology 12 (2022) 972383.

[70] D. Gate, N. Saligrama, O. Leventhal, A. C. Yang, M. S. Unger, J. Middeldorp, K. Chen, B. Lehallier, D. Channappa, M. B. De Los Santos, et al., Clonally expanded cd8 t cells patrol the cerebrospinal fluid in alzheimer’s disease, Nature 577 (7790) (2020) 399–404.

[71] M. E. Pipkin, J. A. Sacks, F. Cruz-Guilloty, M. G. Lichtenheld, M. J. Bevan, A. Rao, Interleukin-2 and inflammation induce distinct transcriptional programs that promote the differentiation of effector cytolytic t cells, Immunity 32 (1) (2010) 79–90.

