## Supplemental results for "Multi-view graph learning for deciphering the dominant cell communication assembly of downstream functional events from single-cell RNA-seq data"

#### 1 Supplementary Table

Table 1: The performance of scDCA for predicting functional states of malignant cells. MSE: Mean Square Error.

| Functional states | Pearson correlation | p-value | MSE |
|---|---|---|---|
| Angiogenesis | 0.683 | <0.05 | 0.037 |
| Apoptosis | 0.879 | <0.05 | 0.010 |
| CellCycle | 0.756 | <0.05 | 0.010 |
| Differentiation | 0.770 | <0.05 | 0.031 |
| DNAdamage | 0.823 | <0.05 | 0.019 |
| DNArepair | 0.875 | <0.05 | 0.009 |
| EMT | 0.670 | <0.05 | 0.032 |
| Hypoxia | 0.816 | <0.05 | 0.024 |
| Inflammation | 0.801 | <0.05 | 0.010 |
| Invasion | 0.776 | <0.05 | 0.027 |
| Metastasis | 0.822 | <0.05 | 0.016 |
| Proliferation | 0.684 | <0.05 | 0.058 |
| Quiescence | 0.736 | <0.05 | 0.028 |
| Stemness | 0.657 | <0.05 | 0.067 |

Table 2: Clinical information of patients in this study.

| Sample | Age at Dx | Sex | ICB |
|---|---|---|---|
| P76 | 68 | F | No ICB |
| P915 | 58 | M | aPD-1 + aCTLA-4 |
| Sample | Duration of ICB Regimen | Time from end of ICB to Biopsy | ICB Response |
| P76 | N/A | N/A | N/A |
| P915 | 2 mo. | 3 mo. | PR |
| Sample | TKI-exposed | Biopsy Site | Histology |
| P76 | No TKI | Kidney | Clear cell |
| P915 | No TKI | Kidney | Clear cell |
| Sample | Grade | Stage | Cell count Post-QC |
| P76 | 4 | IV | 7912 |
| P915 | 3 | IV | 6541 |

TKI: tyrosine kinase inhibitors; PR: partial response.

### 2 Supplementary Figure

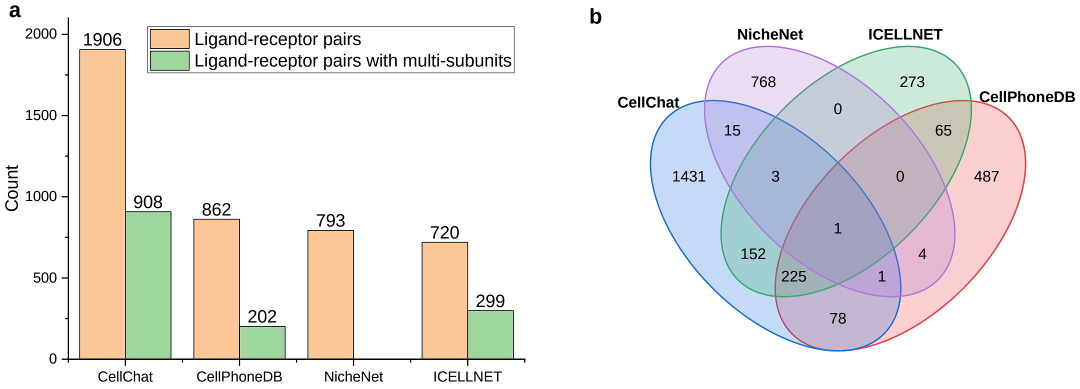


Figure 1: Comparison of the counts of inferred L-R interactions in P76_RNA dataset by CellChat, CellPhoneDB, NicheNet, and ICELLNET, respectively. **(a)** Count of L-R pairs inferred by each of the four tools (NicheNet does not consider the interactions with multi-subunits). **(b)** The overlap between the results of the four tools.

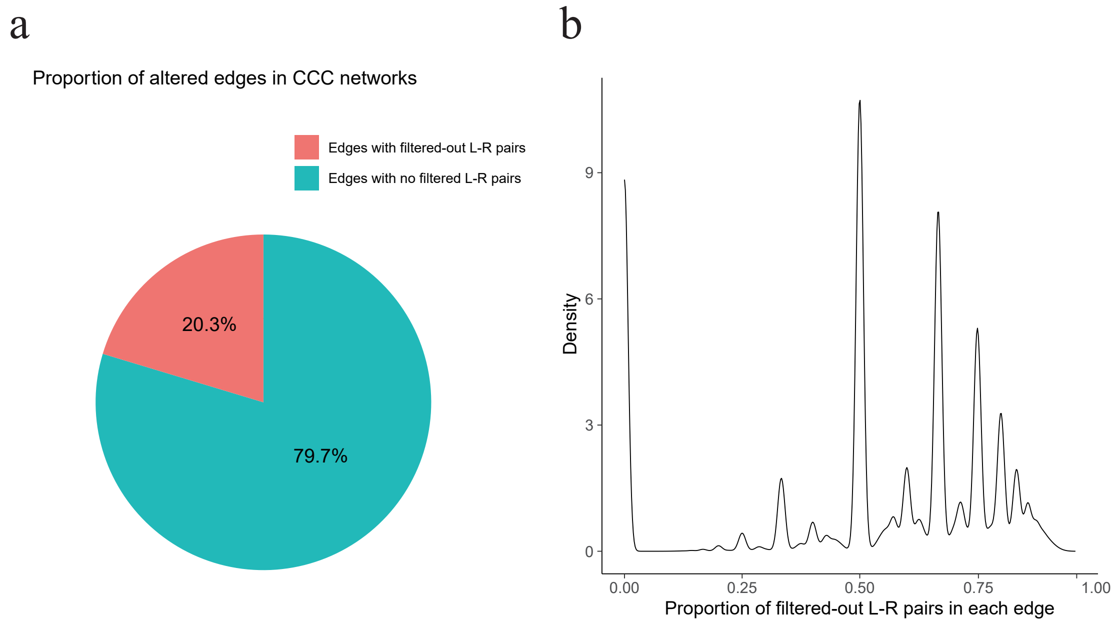


Figure 2: Proportion of the altered edges in CCC network after filtering out L-R pairs with low specificity for P76_scRNA dataset. **(a)** Proportion of edges with filtered-out L-R pairs. **(b)** Proportion of filtered-out L-R pairs on each edge.

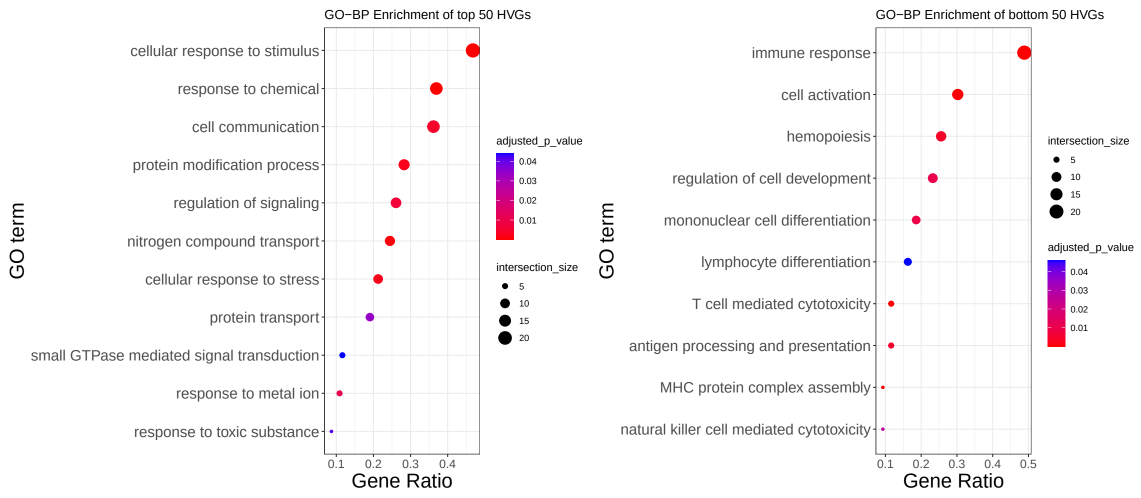


Figure 3: Functional enrichment analysis of GO-BP for the top (left) and bottom (right) 50 highly variable genes (HVGs) with the highest and lowest ratio of $\Delta E$ to total expression levels ($\Delta E + E_0$).

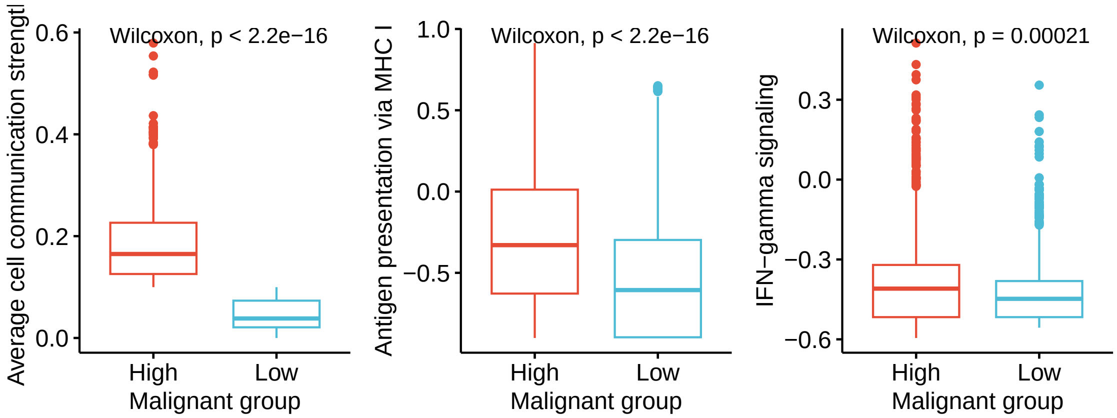


Figure 4: Comparison of the average communication strength with CD8+ T cells, antigen presentation activity via MHC-I molecular, and IFN-$\gamma$ signaling between two groups of malignant cells, which were divided based on their average communication strength with CD8+ T cells.

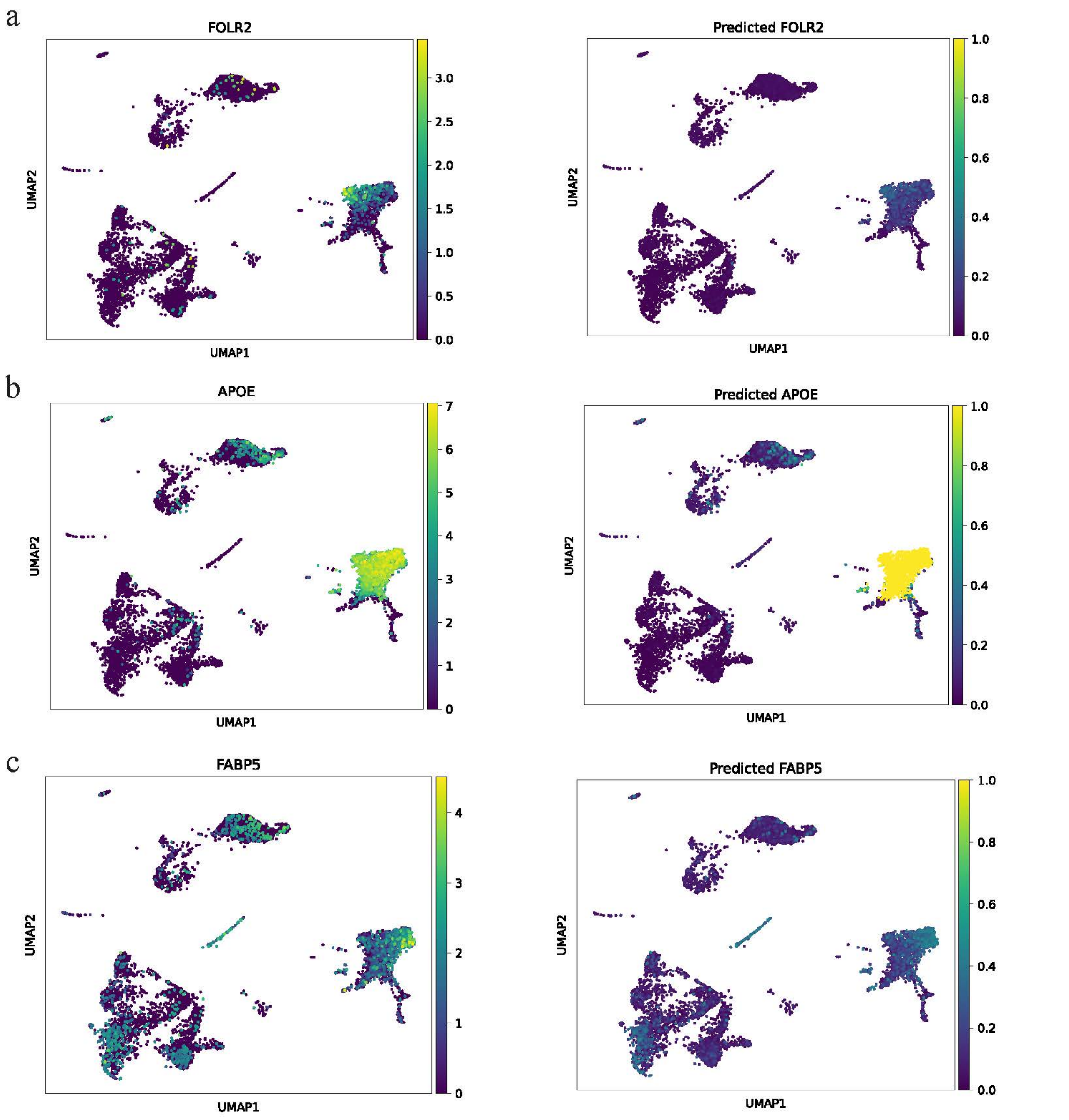


Figure 5: UMAP plots of original expression profiles and scDCA-based predicted expression profiles of three specific genes (FOLR2, APOE, FABP5) in single-cell space.

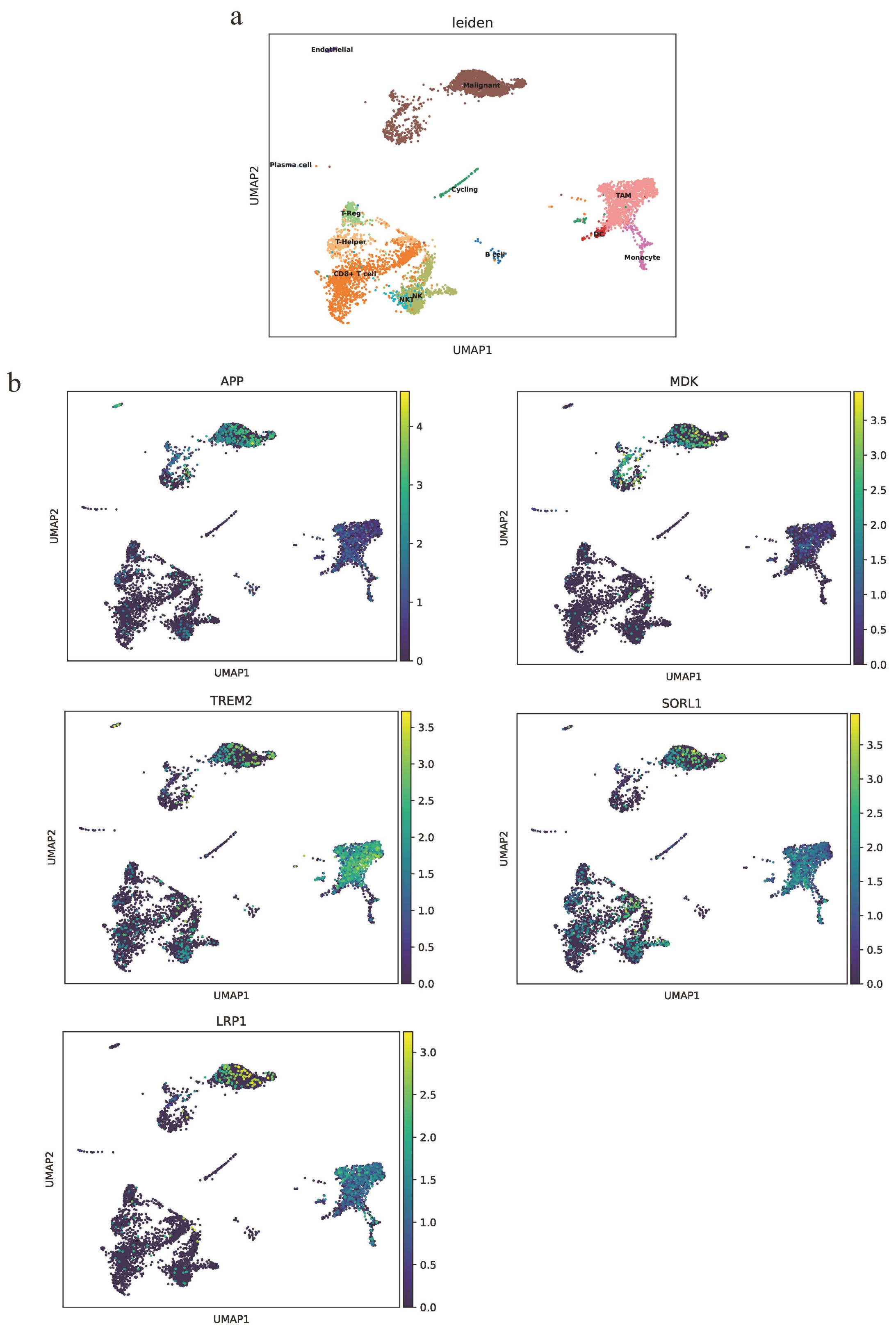


Figure 6: UMAP plots of key ligands and receptors. **(a)** UMAP of all captured cell samples in the data set. Cell type annotation was inherited from the data resource analysis and visualized with different colors. **(b)** UMAP plots of key ligands and receptors between Malignant cells and TAMs that mainly affect FOLR2 expression in TAMs.

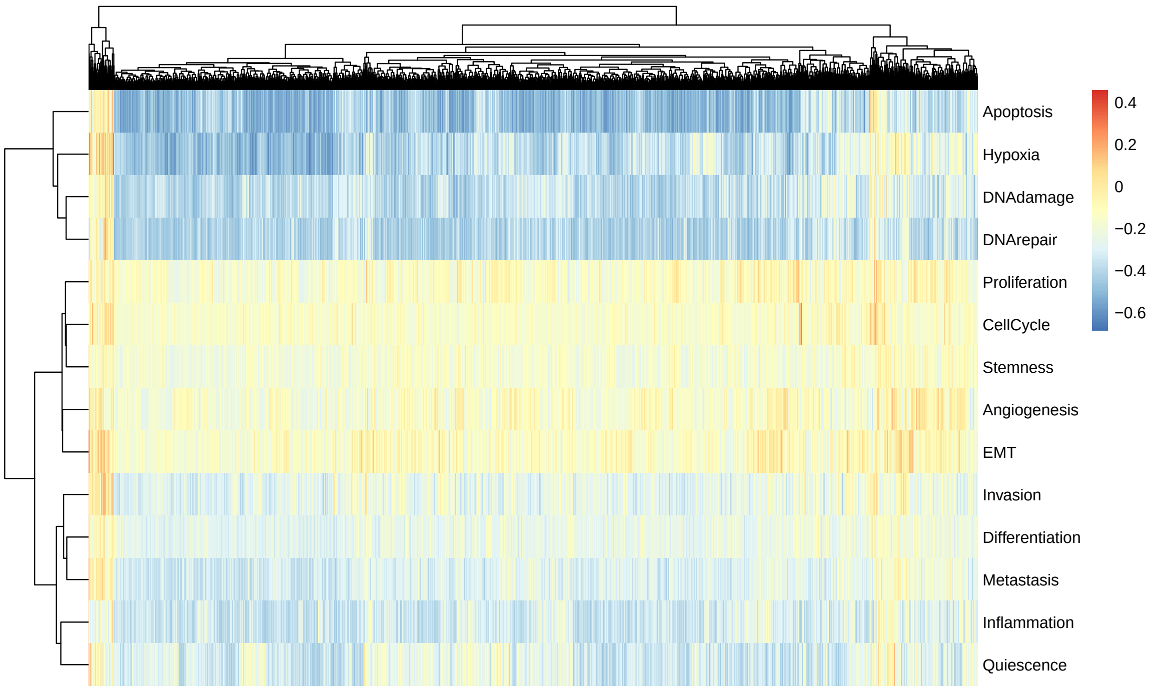


Figure 7: Heatmap showed the activity score of 14 functional states in 7393 tumor cells of patient P76.

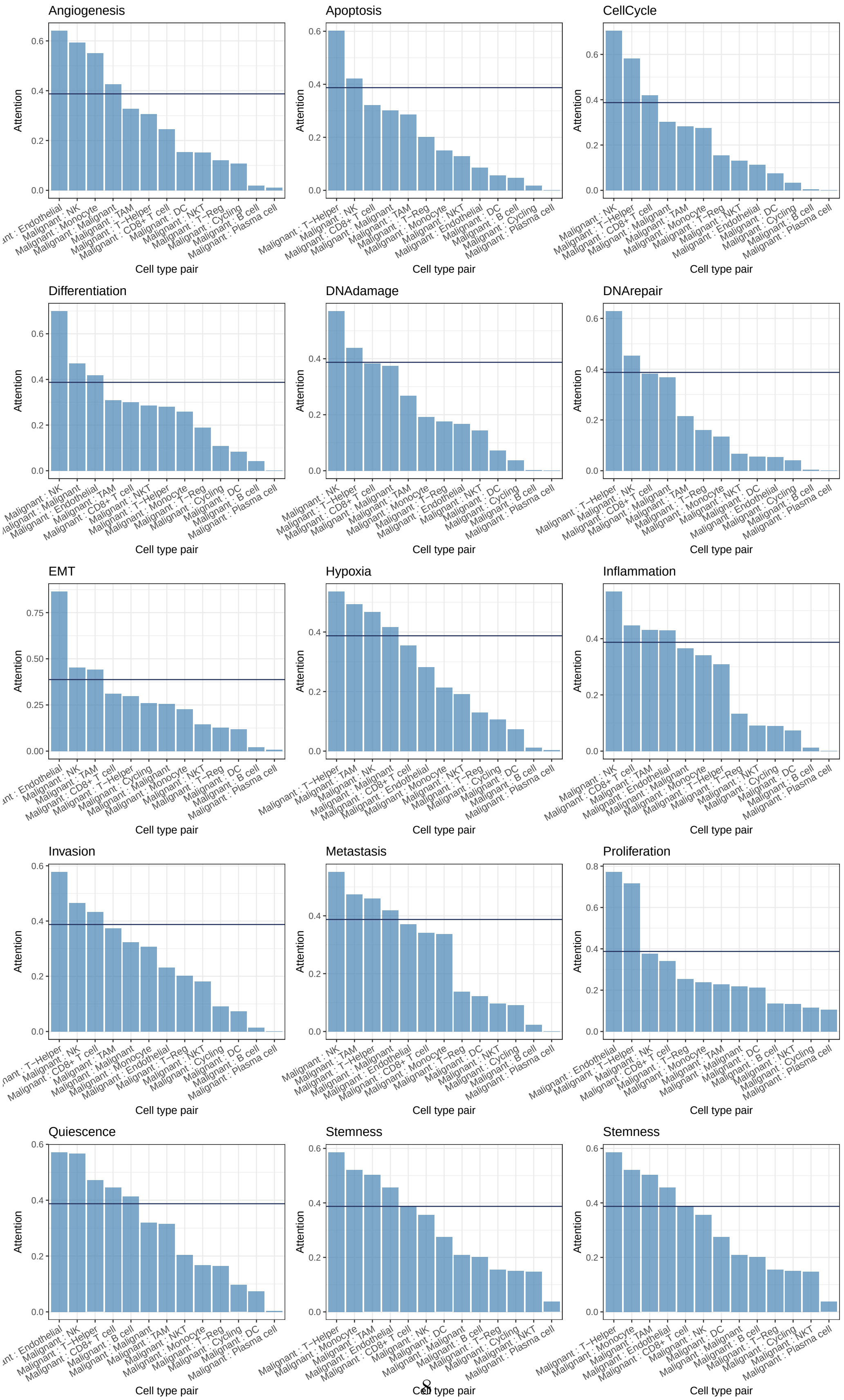


Figure 8: Rank of the attention weight of each cell type pair (x-axis) for each functional state of malignant cells.

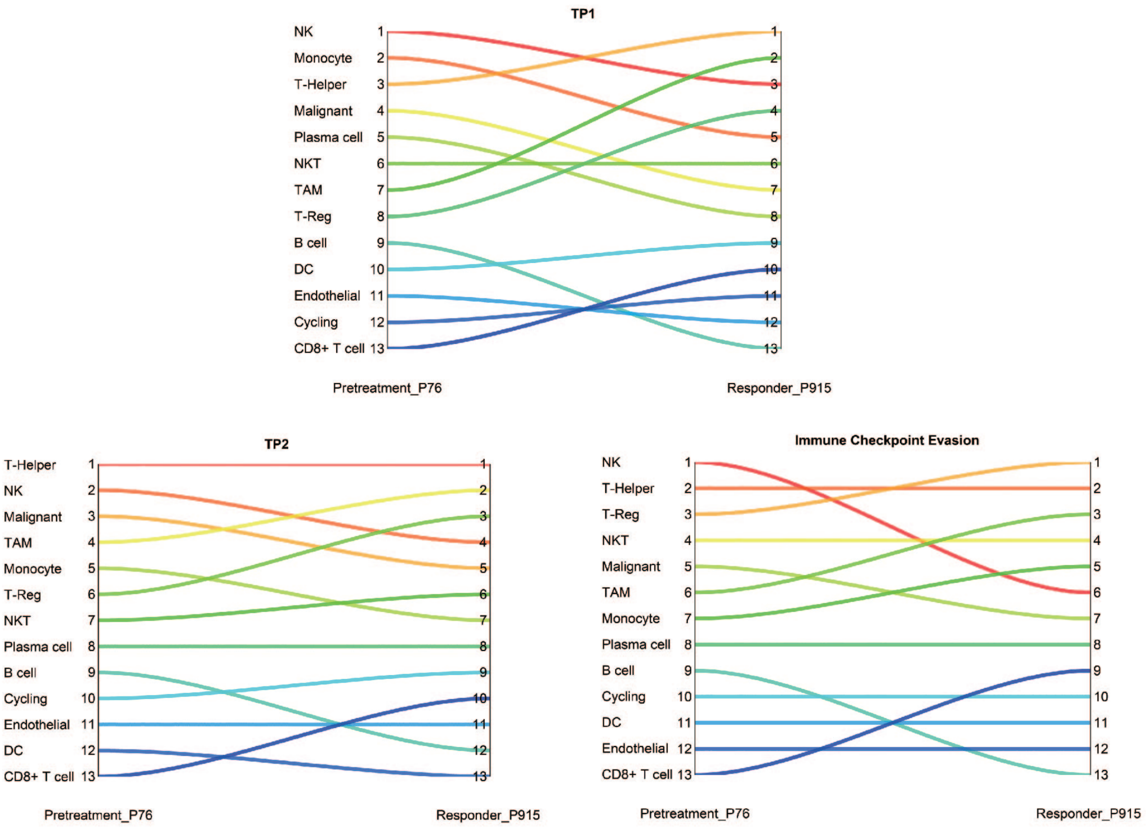


Figure 9: Mapping of the rank changes of DCA for 3 cellular signatures associated with the clinical response of ICB from comparing ICB-responder (P915) to untreated patients (P76).

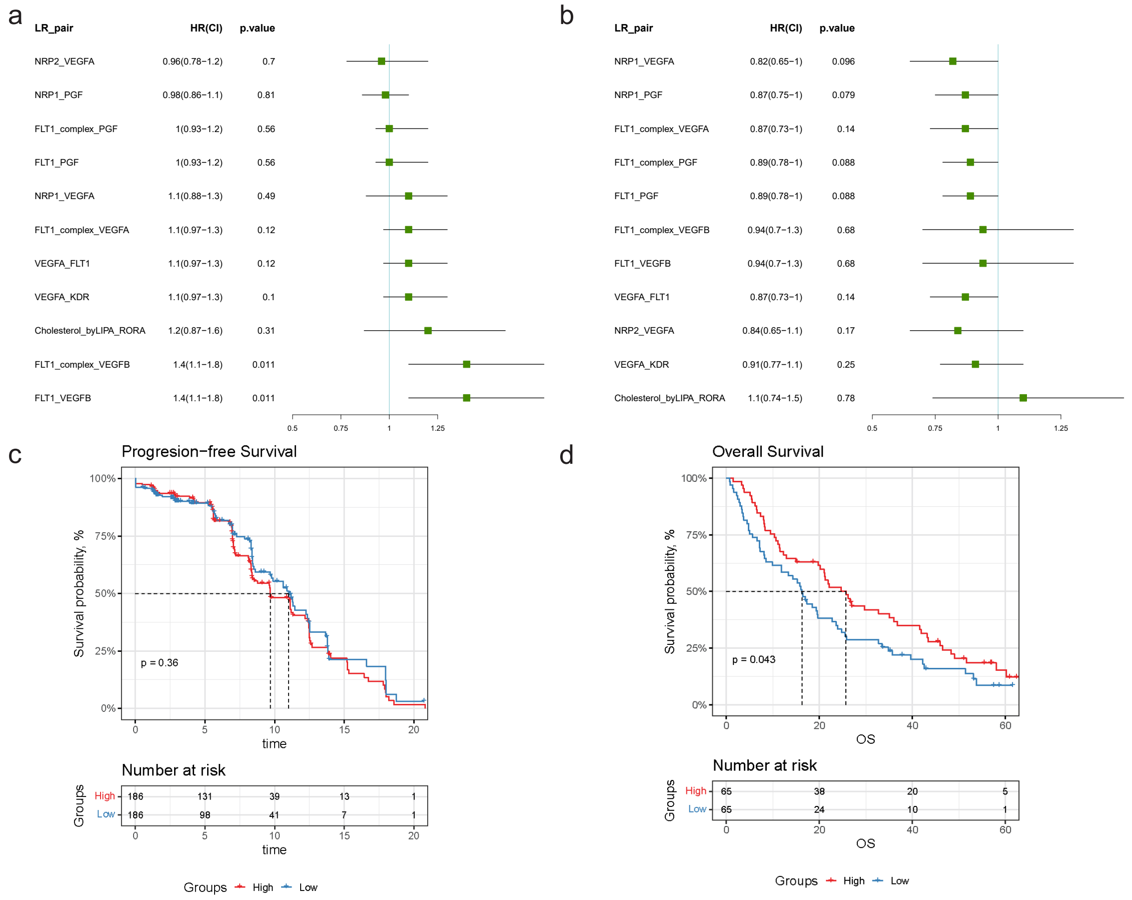


Figure 10: Survival analysis of the integration of the 11 L-R pairs in the two independent cohorts of chemotherapy-treatment-only subpopulations. **(a)** Forest plot of the 11 L-R pairs that exhibited significant prognostic significance (univariate Cox regression analysis) in ICB-treatment-only ccRCC cohorts but not in patients received Sunitinib **(a)** and mTOR inhibitor everolimus **(b)**. Forest plot of the 11 L-R pairs that exhibited significant prognostic significance (univariate Cox regression analysis) in ICB-treatment-only ccRCC cohorts but not in patients who received Sunitinib **(a)** or mTOR inhibitor everolimus **(b)**. Kaplan-Meier curves and log-rank test for the integration of the 11 L-R pairs in the two ccRCC cohorts treatment with Sunitinib **(c)** or mTOR inhibitor everolimus **(d)**.
